# Ribosomal Architecture and rRNA Modification Landscape in the Tick-Borne Parasite *Babesia divergens*

**DOI:** 10.1101/2025.11.11.687446

**Authors:** Cristina Gutierrez-Vargas, Lee S. Izhaki-Tavor, Diana Calvopina-Chavez, Caroline D. Keroack, Pablo Copello, Manoj T. Duraisingh, Mélissa Léger-Abraham

**Affiliations:** Department of Microbiology, Harvard Medical School | Blavatnik Institute, Boston, MA, 02115, USA; Department of Immunology and Infectious Diseases, Harvard T. H. Chan School of Public Health, Boston, MA, 02115, USA; Department of Molecular Microbiology and Immunology, Brown University, Providence, RI 02912, USA; Division of Molecular Medicine, Boston Children’s Hospital, Boston, MA, 02115, USA

**Keywords:** ribosome, cryo-electron microscopy, RNA modifications, Apicomplexan, *Babesia divergens*

## Abstract

*Babesia* is a tick-borne intracellular apicomplexan parasite responsible for diseases ranging from mild to fatal, with a broadening geographic distribution. Due to the complex life cycle of *Babesia* species, their survival depends on the precise control of gene expression, which is primarily regulated by epigenetic, transcriptional, and post-transcriptional mechanisms. High-resolution structural information on key components of the translation machinery, such as ribosomes, could aid in the development of antiparasitic drugs. Here, we report cryo-EM ribosome structures (2.6 Å) from the tick-borne apicomplexan pathogen *Babesia divergens*, showing associated tRNAs, an mRNA fragment, and RACK1, a signaling scaffold crucial to translation regulation. Density map analysis displays ribosome regions at atomic resolution (1.7 Å), which, when combined with nanopore sequencing, enabled the comprehensive identification of rRNA modifications, including modifications unreported in other organisms. The new rRNA modifications localize not only to the reduced *Babesia* rRNA expansion segments but also to functionally essential ribosomal sites, uncovering new avenues for therapeutic intervention against babesiosis.

## Introduction

*Babesia* species are tick-borne apicomplexan parasites that infect a broad spectrum of wild and domestic animals worldwide, resulting in significant economic losses to the livestock industry^1,2^. Among the six *Babesia* species pathogenic to humans, *B. microti* (in the United States) and *B. divergens* (in Europe) are the most prevalent, causing babesiosis^3–6^. In humans, *Babesia* parasites infect erythrocytes, causing malaria-like symptoms^7,8^. Treatment typically involves a combination of antibiotics and antimalarials, including azithromycin with atovaquone or clindamycin with quinine^9^. Babesiosis can also be fatal, especially in immunocompromised individuals, in whom prolonged treatment, sometimes extending over two years, is often required to clear the infection^10–12^. Drug pressure increases the risk of resistance mutations, potentially leading to chronic infection or relapsing disease^13^. The development of new therapeutics targeting *Babesia*-specific molecular pathways, rather than relying on repurposed drugs (e.g., combination of antibiotics and antimalarial drugs, such as azithromycin and atovaquone or clindamycin and quinine), is urgently needed. This need is further underscored by the expanding geographic distribution of babesiosis and its co-infection risks in northern regions, particularly in the mid-Atlantic region of the USA^14,15^.

Apicomplexan parasites, such as *Plasmodium*, *Toxoplasma,* and *Babesia*, have complex life cycles^16–18^. Their survival depends on the precise control of gene expression, which is primarily regulated by epigenetic, transcriptional, and post-transcriptional mechanisms^19,20^. As they transition between arthropod vectors and mammalian hosts and progress through the various stages of their life cycle, translation regulation becomes crucial so that parasites can quickly adapt their protein expression level accordingly. Nonetheless, the mechanisms underlying protein synthesis in apicomplexan parasites remain understudied compared to those in bacteria, yeast, or humans.

Protozoan ribosomes present unique regulation opportunities, as they exhibit heterogeneity and plasticity in their ribosomal protein composition, recruitment of accessory proteins, and the types of ribosomal RNA (rRNA) incorporated^21^. Ribosomal RNA modifications are also of particular interest because they have been linked to altered translation landscapes with biological relevance^22–24^; for example, they differ in patterns between healthy and tumor cells^25–27^, or by altering antibiotic binding in bacteria as a mechanism of antibiotic resistance^28^. Although only a few rRNA modifications have been reported in a recent cryo-EM ribosome structure from *P. falciparum*^29^, no comprehensive characterization of rRNA modifications has been conducted concomitantly with ribosome structural analysis from any apicomplexans^29–32^, limiting our understanding of the mechanisms and potential roles of individual modified nucleotides in these parasites.

We report here an atomic model of the cytoplasmic ribosome from the tick-borne apicomplexan pathogen, *Babesia divergens*, at a global resolution of 2.6 Å, providing an unparalleled view of both the ribosomal protein and rRNA architecture^29,30,32–35^. The model shows bound tRNAs associated with an mRNA fragment and the regulatory scaffold protein RACK1, which is integral to translation control. Using high-resolution cryo-EM maps in combination with nanopore sequencing, we thoroughly characterize rRNA modifications and identified modification positions not previously observed in any other organism. These novel modifications are present not only within the minimized rRNA expansion segments of *Babesia* but also at key functional sites of the ribosome, highlighting novel avenues for therapeutic intervention against babesiosis.

## Results

### Overview of the *Babesia* ribosome high-resolution structure

We isolated *Babesia* ribosomes from an asynchronously grown *Babesia divergens* Rouen 1987 strain. Following ribosome purification, cryo-EM sample preparation, and image processing, including particle sorting, we obtained two main conformational states that had the typical and distinctive morphology of 80S ribosomes (**Figs. S1 and S2**, see *Methods*). In one conformational state (3D class 1), the ribosome particles contain three tRNAs occupying the E-, P-, and A-sites, and an mRNA fragment (approximately nine nucleotides) encompassing the tRNA binding sites (**Fig. 1a, left panel and Table S1**). Consistent with this conformational state, the two adenosines within helix 44 (small subunit h44), responsible for monitoring the first two base-pairs of the codon-anticodon mini helix (A1683 and A1684, corresponding to human A1824 and A1825) are in a flipped-out conformation, stabilizing and favoring the accommodation of the A-site tRNA in the ribosome during the tRNA selection step (**Fig. 1a, right panel**). In the second conformational state (3D class 3), the ribosome particles contain an E-site tRNA (**Fig. 1b and Table S2**), where the A1683 remains stacked within h44 (data not shown). The corresponding density for A1684 is unresolved, consistent with increased flexibility when the A-site is unoccupied. We used local CTF refinement, 3D re-refinement, and post-processing in RELION for both conformational states, followed by masking and multibody refinement. For the three tRNAs-bound ribosome conformational state, we obtained final 80S maps with a global resolution of 3.1 Å. For the E-site tRNA-bound ribosome, the global resolution is 2.6 Å. Although we built atomic models for both conformational states (ribosomes with three tRNAs and mRNA versus ribosomes with only an E-site tRNA), all our analysis originates from ribosomes bound to the E-site tRNA, due to its higher resolution, with areas within the ribosome core reaching atomic resolution (approximate resolution of 1.7 Å, **Fig. 1c**).

**Fig. 1.**
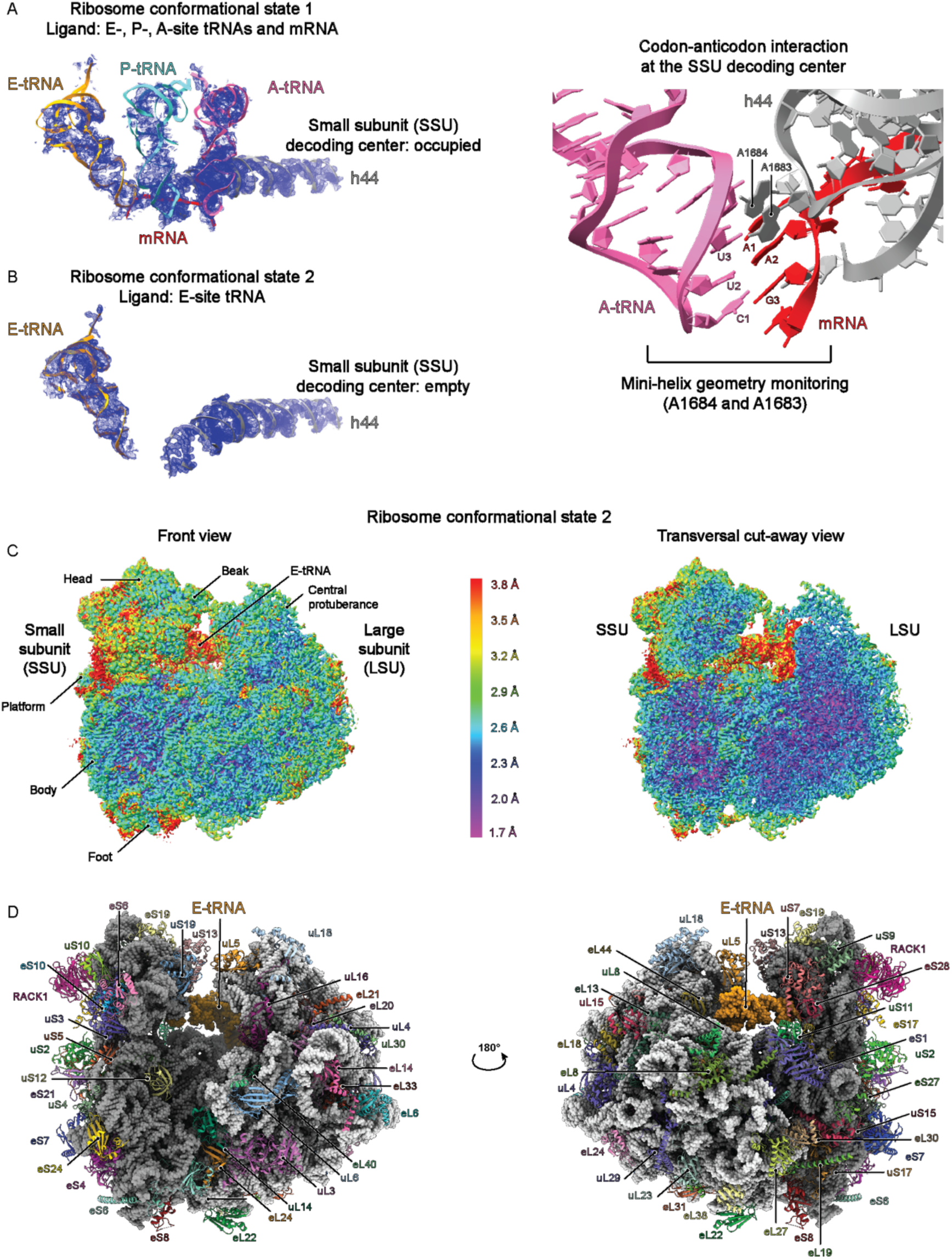
Overview of Babesia ribosome structures. **A** Left panel: Partial density from the consensus maps corresponding to the three tRNAs (E-, P-, and A-site tRNAs) and an mRNA fragment bound to the ribosome (Conformational state 1, 3D Class1), see **Figs. S1, S2, and Table S1**). Right panel: Enlarged view of the decoding center. A1683 and A1684 are in a flipped-out conformation relative to helix 44 (h44) of the small subunit (SSU) and monitor the geometry of the first two base-pairs of the codon-anticodon mini-helix. **B** As described in ***A***, but for the conformational state 2 (3D Class 3), where a single E-tRNA is bound to the ribosome, and that is further described in **C** and ***D*** (see **Figs. S1, S2, and Table S2**). **C** Left panel: Sharpened multi-body refinement maps of the *Babesia* ribosome bound to an E-site tRNA. The combined individual bodies are colored according to the estimated local resolution (color code in the middle). The morphological features of both subunits are labeled. LSU: large subunit. Right panel: Transversal cut-away view showing the highest resolution (1.7 Å) within the ribosome core **D** Left panel: Ribosome model for the corresponding cryo-EM maps shown in ***c***. The ribosomal proteins are shown in ribbon colored representations. The rRNA is represented in dark and light gray spheres for the SSU and the LSU, respectively, and the E-site tRNA is in orange. Right panel: 180-degree rotation along the Y axis.

To build the ribosome protein component of the atomic model (see *Methods*), we initially used a blast search originating from *Plasmodium falciparum* 3D7 (PDB IDs: 3J7A, 3J79)^30^ to *Babesia divergens* 1802A strain (**Tables S3 and S4**). The identified sequences were then implemented in Alphafold2^36^ to obtain structure prediction, which were then docked into the *Babesia* maps using ChimeraX^37^. We also analyzed and confirmed the complement of ribosomal proteins from purified ribosome samples by LC/MS-MS (**Tables S5-S7**). For the rRNA component, we used sequences retrieved from RNAcentral^38^, generated sequence alignment between *Babesia* and *Toxoplasma* using Clustal Omega^39^ and obtained secondary structure prediction maps using RNAcentral. We selected conserved fragments to initiate model building. Iterative rounds of manual model adjustment (Coot)^40^ and model refinement (PHENIX)^41^ led to the final atomic model of the *Babesia* cryo-EM ribosome structure (**Fig. 1d, Tables S1 and S2**). The *Babesia* atomic model has a higher resolution than any other apicomplexan ribosome structure previously determined^29,30,33,34,42^, as reflected in the better-defined cryo-EM maps, especially in the small subunit (SSU) (**Fig. 2a-c**).

**Fig. 2.**
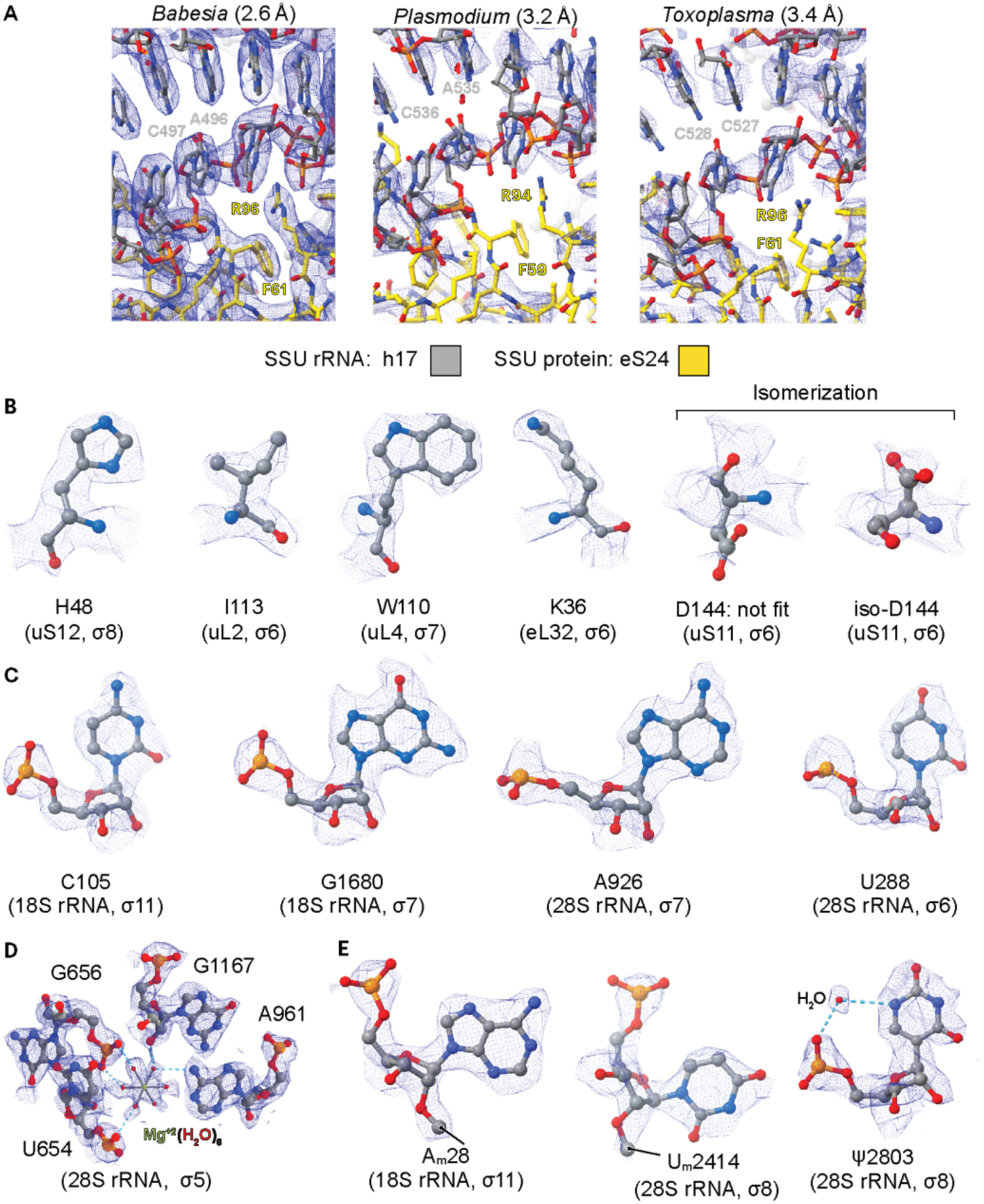
High-resolution details of the *Babesia* ribosome bound to an E-site tRNA. **A** Global views of the SSU estimated with RELION (FSC=0.143) centered on the body (h17: C497 and A496 in gray, and eS24: R96 and F61 in yellow), illustrating the quality of the cryo-EM maps presented in this study when compared to those of *P. falciparum* (EMD-2660, PDB 3J7A, threshold level of 0.2) and *T. gondii* (EMD-6780, PDB 5XXU, threshold level of 0.06). **B** An atomic view of residues from the SSU and LSU, where the side chains are clearly defined for different residue types, including the fitting of Asp144 isomerization from uS11. **C** Examples of nucleotides from the 18S (SSU) and 28S (LSU) rRNA, where the ring structures of the bases and riboses are well resolved. **D** Atomic view of the interaction between the H25.1 loop from the LSU with a magnesium ion (green). Gray dashed lines correspond to the first-shell interactions between water molecules (red spheres) and the magnesium ion, while cyan dashed lines indicate hydrogen bonds formed between water molecules and rRNA nucleotides. **E** An atomic view of a few nucleotide modifications observed throughout the *Babesia* ribosome structure identified by cryo-EM density analysis, such as methylations (indicated by a line) and pseudouridylation (coordinated by a water molecule). Nanopore sequencing experiments confirmed the methylation and pseudouridylation of A28, U2414, and U2803, respectively (see *Methods)*.

Our structure includes water molecules and ions (Zn^+2^ and Mg^+2^, **Fig. 2d**), which were assigned based on their typical chemical coordination characteristics (e.g., tetrahedral vs octahedral), coupled to the strength and quality of the density maps within specific areas. We also observed post-transcriptional modifications (see *below*), such as rRNA modifications (**Fig. 2e**), as well as post-translational modifications, including residue isomerization (**Fig. 2b**), which was also recently described in human ribosomes^43^.

### Features of the *Babesia* ribosomal proteins

We modeled more than 94 % of the ribosomal protein residues within the *Babesia* ribosome atomic model, identifying 33 and 41 ribosomal proteins for the SSU and LSU, respectively (**Tables S3 and S4**). We found that *Babesia* uS7 (residues 70-97), eS10 (residues 26-46), uS12 (residues 64-106) (**Fig. S3**), eL18 (residues 122-146) and eL24 (residues 61-72) (**Fig. S4**) lack these sequence regions, as initially annotated in the *Babesia* 1802A strain genome. These findings are based on the density map analysis, which shows no corresponding density for these portions of the protein sequences. RNA-seq data also supports these observations, indicating the presence of introns that would remove the sequences coding for each of those regions^44^.

We did not identify an eS17 protein sequence in *B. divergens* 1802 using the *P. falciparum* sequence (PF3D7_1242700) in a blast search, but did identify a corresponding sequence in *B. bovis* (BBOV_III001250). Genomic analysis using *B. bovis*’s sequence led us to identify a corresponding protein sequence in *B. divergens* Rouen 1987 strain at the CCSG02000001.1: 1890202..1890772 supercontig, for which there is a genome sequence gap in the *B. divergens* 1802 reference strain. The eS17 *B. divergens* sequence differs from the *B. bovis* sequence at four positions, which were modeled accordingly in the atomic model (*B. bovis* numbering: R3G, A22G, S25G, and Q40L) (**Fig. S5a**). A similar strategy, using *T. gondii* as a reference sequence (TGME49_213580), led us to identify the unannotated genomic sequence corresponding to eL41 in the *B. divergens* 1802 reference strain (JAHBMH010000034: 30674..30988 supercontig). The protein sequence matched the density maps, including the N-terminal sequence of eL41 (residues 1-14), which is absent in humans but present in *P. falciparum*, *T. gondii*, and other *Babesia* species (**Fig. S5b**). We did not model the proteins forming the ribosome L1 (uL1) and P stalks (uL10, uL11, P1, P2), because their corresponding density were too weak, despite being present in our sample based on mass spectrometry analysis (**Tables S5-S7**). Both stalks are known to be flexible, with the L1 stack binding the deacylated-tRNA during translocation and regulating its release, while the P-stalk helps recruiting and stimulating the ribosome-associated GTPase translation factors^45–48^.

### Features of *Babesia* RACK1

In addition to the ribosomal protein complement, the eukaryotic ribosome binds to the receptor for activated C kinase 1 (RACK1)^49–51^, a scaffolding and signaling protein^52^ that increases the translation efficiency of capped mRNA and contributes to heterogeneity and diversification within the ribosome population^21^. RACK1 has been implicated in various diseases, including cancer progression, inflammation, and neurological disorders^51^. RACK1 interacts with the SSU head domain near the mRNA exit tunnel (**Figs. 1d, 3, and S6**)^50^. In the human ribosome, the C-terminal tail of uS3 mediates contact with the propeller blades 4 and 5 of RACK1 (**Fig. S7**). At the same time, RACK1 interacts mainly with eS17 and uS9, and with h39 and h40 through a network of electrostatic interactions (Arg/Lys patch, **Figs. 3 and S6**)^43,49,53^. The molecular details of RACK1 and ribosomes purified from apicomplexans untreated with any drugs have not been reported, as density for RACK1 was either not observed or very weak, and unmodeled in *Plasmodium* and *Toxoplasma* single-particle cryo-EM ribosome structures, respectively^29,30,33,34^. In apicomplexans, including *Babesia*, the C-terminal tail of uS3 is significantly shorter than in humans (**Fig. 3d**). It was thus suggested that this loss of contacts would substantially destabilize the RACK1/ribosome interactions in apicomplexans. The *Babesia* ribosome structure presented in this study includes RACK1, for which the apparent density corresponds to a high level of occupancy on the ribosome. Our atomic model lacks the last ten residues from the C-terminus of uS3, but based on the human ribosome structure comprising RACK1, four additional residues would be required to reach the propeller blades of RACK1. Therefore, even if uS3 were fully modeled, uS3 would not reach RACK1. Although the interactions between RACK1 and eS17 are of the same nature in human ribosomes and in the *Babesia* ribosome (hydrogen bonds, salt bridges, and hydrophobic contacts), the binding interface is larger in *Babesia*, where five eS17 residues interact with RACK1 compared to four in humans. This observation would not be sufficient to explain the absence of RACK1 in the other apicomplexan ribosome structures mentioned above. Other factors may contribute to this discrepancy, such as during cryo-EM sample preparation or the fact that the *Babesia* ribosome preparation encompasses all the host-parasite’s life stages, as the culture was asynchronously grown.

**Fig. 3.**
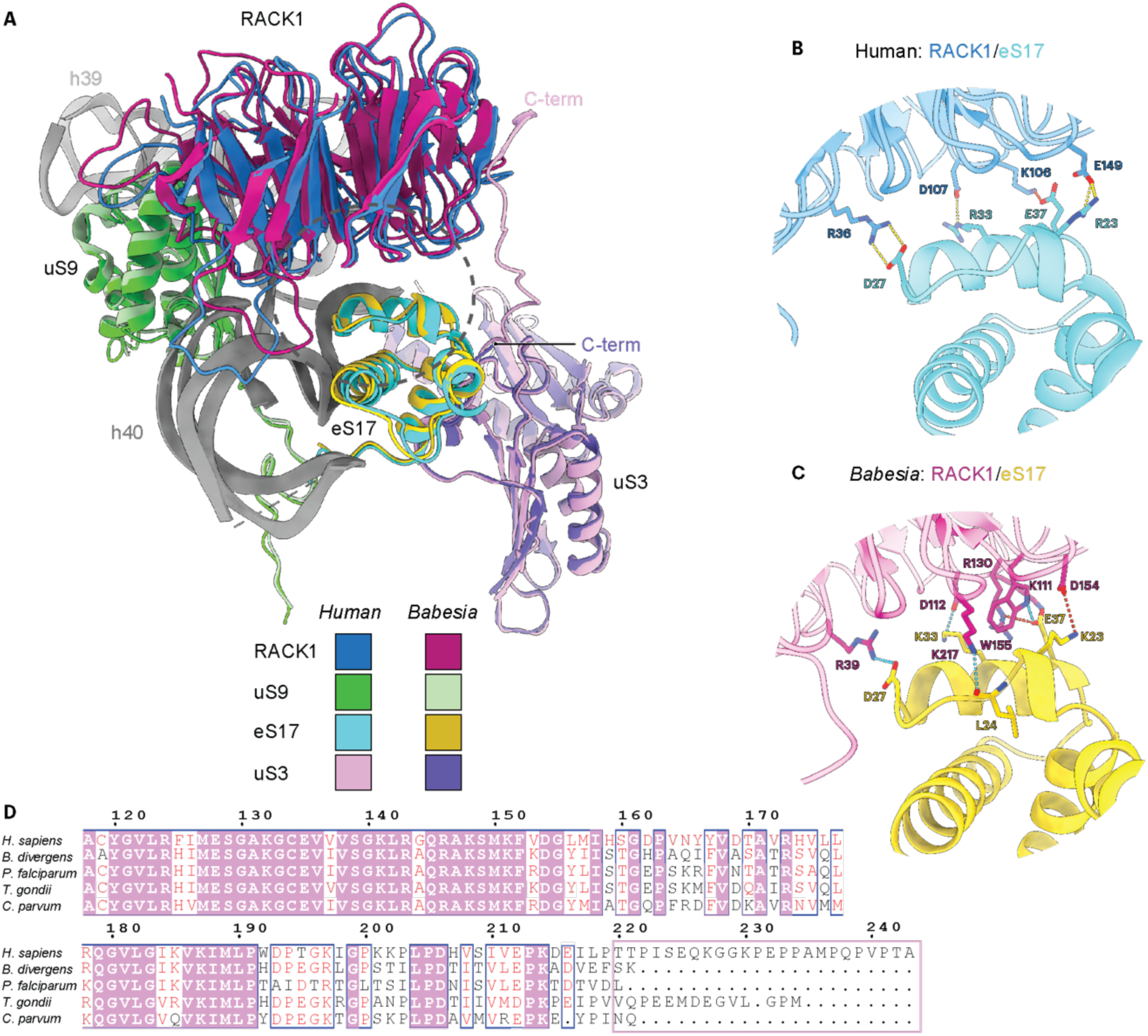
Molecular details of the interaction between RACK1 and eS17. **A** An overview of the localization of RACK1 within the head of the SSU, where it is surrounded by uS3, uS9, eS17, and helices 39 and 40 (see also **Figure 1).** Human (PDB 8QOI) and *Babesia* proteins are colored and labeled accordingly, with their respective C-terminus indicated (C-term). A dotted gray circle corresponds to the zoomed region shown in panels ***B*** and **C. B** Key interactions between RACK1 and eS17 from humans. The hydrogen bonds and salt bridges are displayed with dotted yellow and red lines, respectively, **c** Key interactions between RACK1 and eS17 from *Babesia.* The hydrogen bonds and salt bridges are displayed with dotted cyan and red lines, respectively. **D** Partial sequence alignment of uS3 showing the sequence divergence at the C-terminus of uS3 (pink rectangle). Sequence alignment was generated using Clustal Omega and displayed using ESPript3. White letters on a pink background correspond to identical residues, while red letters on a white background indicate conservation. The numbering is based on the human sequence. VEuPathDB gene IDs: *H. sapiens:* P23396, *B. divergens:* Bdiv_0H570, *P. falciparum:* PF3D7_1465900, *T. gondii:* TGME49_232300, *C. parvum:* cgd7_2250.

### Features of the *Babesia* rRNA

The secondary structure of rRNA is highly conserved across all domains of life. Eukaryotes have acquired multi-nucleotide insertions, termed expansion segments (ESs), that emerge from the universal rRNA core relative to prokaryotes. These ESs are mostly located on the ribosome’s surface, distant from the conserved functional centers, and exhibit remarkable variability in both sequence and length across species. While the functions of these ESs remain poorly understood, several studies have reported implications in ribosome biogenesis^54–56^, translation regulation^57^, and in the fidelity and accuracy of protein synthesis^58,59^. We thus compared human ESs with those of *Babesia* to identify differences that could further our understanding of their roles across evolution.

The human SSU ESs are all found in *B. divergens* (**Table 1**). While most are conserved, with less than 20 % sequence length variation, a few are truncated (ES3, ES6, and ES12), all located at the solvent-exposed side of the SSU. No function has been attributed to these ESs thus far. The ESs from the LSU exhibit significantly greater variability than those from the SSU. For example, the higher-eukaryote (vertebrate) specific segments (ES30L and ES44L) are absent from *Babesia*, many ESs are substantially truncated, and several ES-specific helices (ES7Ld-h, ES9La, and ES20La) are missing.

**Table 1.**
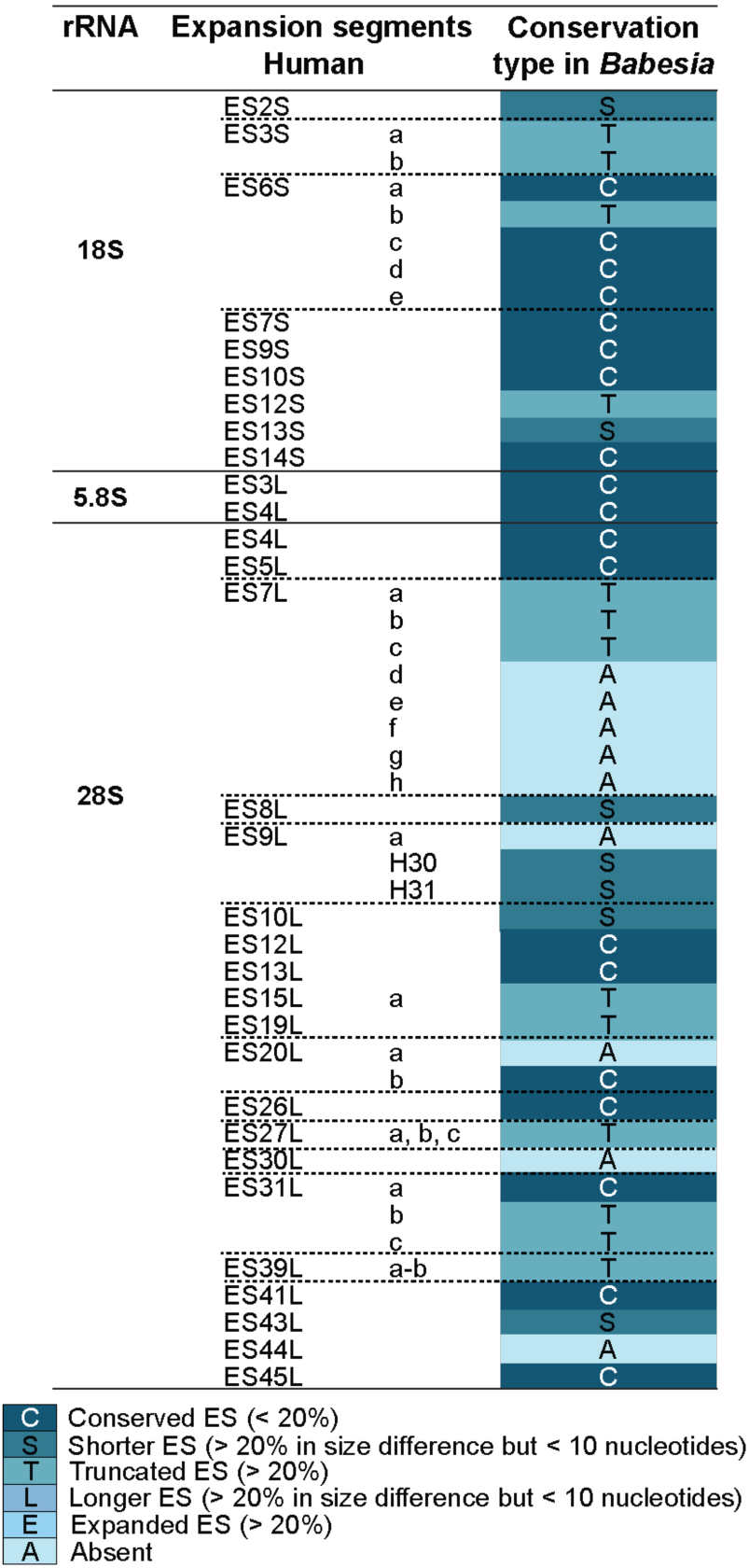
Conservation degree analysis of *Babesia* ESs when compared to human Ess.

*Babesia* ESs are generally conserved or truncated compared to *T. gondii* and *P. falciparum*, respectively (**Table 2**)^30,34^. Despite a closer phylogenetic relationship to *Plasmodium*, *Babesia* ES lengths are more comparable to those of *Toxoplasma*. This is likely due to lineage-specific expansion of rRNA in *Plasmodium* species, which contains several species-specific insertions (ES7LB1, ES34L, ES36L) and unusually long ESs (ES6S, ES7S, ES9S, ES10S). In contrast, the rRNA of *Babesia* is notably compact, consistent with its reduced genome size^60–62^.

**Table 2.**
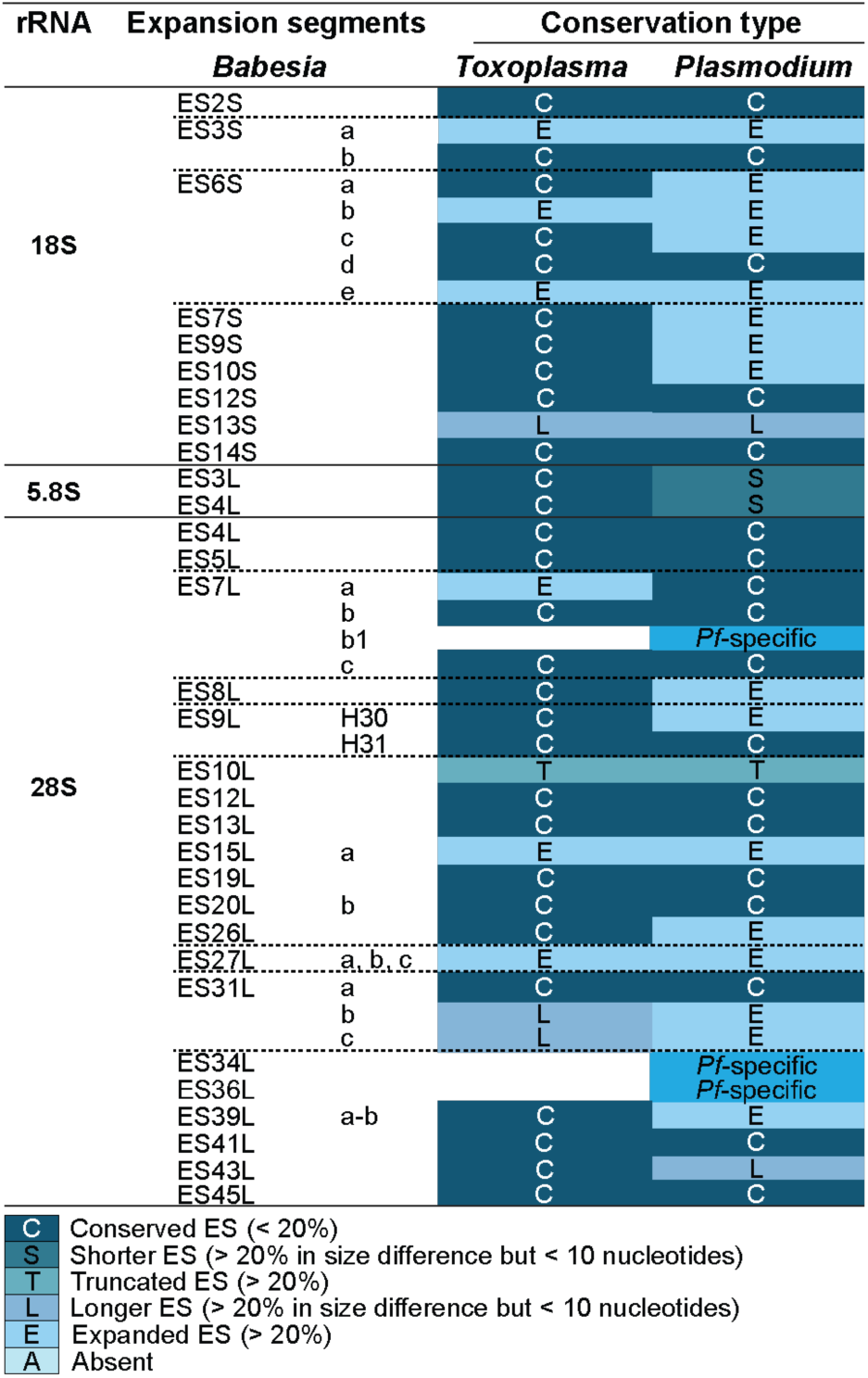
Conservation degree analysis of *Babesia* Ess when compared to *Toxoplasma* and *Plasmodium* ESs.

Much of the structural divergence between *Babesia* and closely related apicomplexans is concentrated in a cluster of expansion segments located at the back of the large ribosomal subunit: ES7L, ES10L, and ES15L (**Fig. 4**). In eukaryotes, ES7L is of particular interest, as it interacts with numerous ribosomal assembly factors, highlighting its significance during ribosome biogenesis^63^. ES7L is the largest eukaryotic expansion segment and is subdivided into eight helices (ES7La-h). In *Babesia* and *Toxoplasma*, ES7L comprises only three helices (ES7La, b, and c), but ES7La is truncated by more than 50 % in *Babesia* relative to *Toxoplasma* (**Fig. 4a and Table 2**). The adjacent *Babesia* ES15L is also markedly truncated when compared to *Plasmodium* and *Toxoplasma* (**Fig. 4a**). While ES7Lb interacts with the ribosomal protein eL14, we were unable to model ES7Lc due to blurred density, indicative of its flexibility (**Fig. 4b**). This is contrasting with the equivalent area in *Toxoplasma*, where ES15L forms a pseudoknot with ES7Lc, which most likely reduces its flexibility (**Figs. 4c and 4d**).

**Fig. 4.**
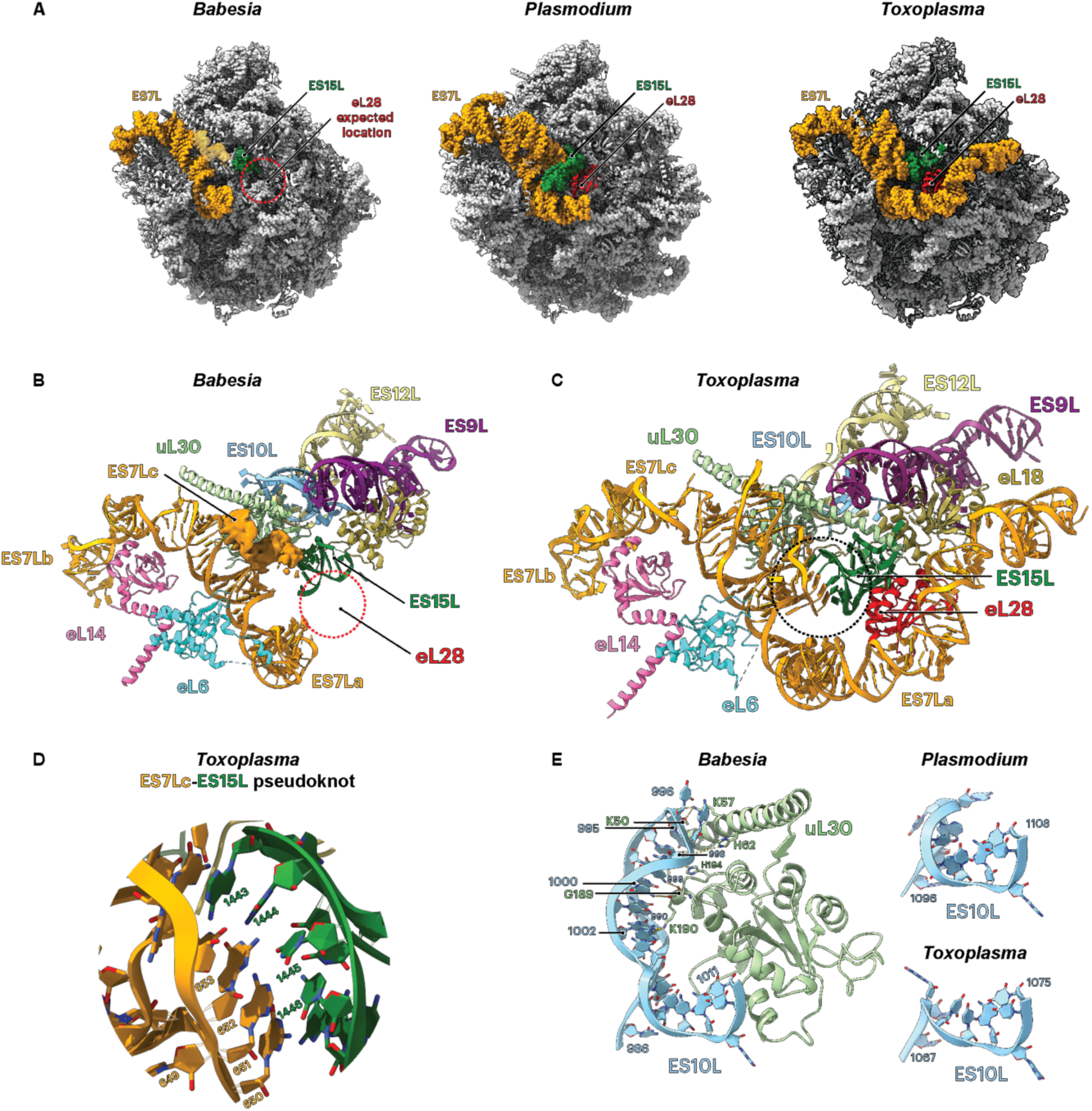
Molecular details of the expansion segments (ES) from the *Babesia* large subunit (LSU) in comparison to *Plasmodium* and *Toxoplasma.* **A** The large ribosomal subunit is shown from the solvent side. General overview of the ESs localization that are mentioned in the text: ES7L and ES15L are labeled and colored, respectively, in orange and green. The ribosomal protein eL28 (red), present in *Plasmodium* and *Toxoplasma,* is absent from the *Babesia* ribosome atomic model (indicated by a dashed red circle). **B** Interaction details within the vicinity of ES7L, ES10L and ES15L in *Babesia,* emphasizing the shorter ES7L and ES15L, the unmodeled ES7Lc, and the absence of eL28. **C** Interaction details within the vicinity of ES7L, eL10L and ES15L in *Toxoplasma,* highliting the longer ES7L RNA fragment. **D** ES7Lc and ES15L form a pseudoknot between nucleotides 650-653 and 1443-1446, which is shown as an enlargement of ***C* E** Comparison of ES10L between *Babesia* and *Plasmodium* and *Toxoplasma* ES10L. Nucleotides from *Babesia* ES10L interacting with uL30 residues are indicated, while the shorter ES10L from *Plasmodium* and *Toxoplasma* do not sustain these interactions.

eL28 is absent from the *Babesia* ribosome cryo-EM density (**Fig. 4a**). Although we identified eL28 homologs in other *Babesia* species that are pathogenic to humans, such as in *B. duncani* and *B.* microti, we did not identify eL28 in the genome of *B. divergens* strains. eL28 was also absent from mass spectrometry data analysis (**Table S6**). Within the same vicinity, ES10L in *Babesia* is expanded compared to *Plasmodium* and *Toxoplasma,* respectively (**Figs. 4b, c and e**). *Toxoplasma* ES10L is found underneath ES9L and ES12L (**Fig. 4c**). In contrast, the *Babesia* extended ES10L tethers through the solvent-exposed surface and contacts predominantly the ribosomal protein uL30 (**Figs. 4b and 4e**). In combination, the remodeling of ES7L, ES10L, and ES15L results in a significant modification of the LSU surface.

### *Babesia* rRNA modifications

Ribosomal RNA modifications are of interest due to their roles in translation regulation and antibiotic resistance^28^. Yet, they have not been comprehensively characterized alongside ribosome structures in apicomplexans, where only a few were reported in a recent *P. falciparum* cryo-EM ribosome structure^29^. We assigned rRNA modifications based on protruding density that is continuous with the ribose or base in the high-resolution maps of the *Babesia* ribosome containing an E-site tRNA (Conformational state 2, 3D class 3). Our search was also cross-referenced with known regions of the ribosome containing rRNA modifications, as described in human ribosomes (near atomic resolution) and in *Leishmania* parasites, where both studies reported a high number of rRNA modifications^24,43,64^. We complemented our strategy using nanopore sequencing^65,66^, which enabled us to identify rRNA modifications in areas of the ribosome associated with lower resolution. Also, because we used asynchronous cultures, nanopore sequencing allows us to identify rRNA modifications that may be present at a specific developmental stage and thus be underrepresented in the dataset. For this reason, we applied two different thresholds during nanopore sequencing analysis: modifications identified at more than 50 % (≥ 50 %), and modifications identified between 30 and 50 % (≥ 30 % and ≤ 50 %) at a given position (see *Methods*).

By combining cryo-EM density map analysis with nanopore sequencing, we identified a total of 66 methylation sites (**Fig. 5**). Out of the 66 methylation sites identified, 59 were detected using nanopore sequencing, where more than 80 % are within the ≥ 50 % threshold. Among those 59 rRNA methylations, about half were new methylation sites, with a quarter of those being observable in the cryo-EM density maps. Among all the methylations identified, 2’-*O*-Me modifications were the most frequent, with very few m^1^ and m^6^ nucleotide methylations. In the SSU, most new methylations are located within the decoding center, the mRNA channel, and the head–body interface that mediates the SSU swiveling along with tRNA dynamics (**Figs. 5a and 5b**). In the LSU, newly reported methylations cluster within the peptidyl transferase center (PTC), including helix 80 (LSU H80), which binds the amino-acid extremity (CCA-end) of the P-site tRNA.

**Fig. 5.**
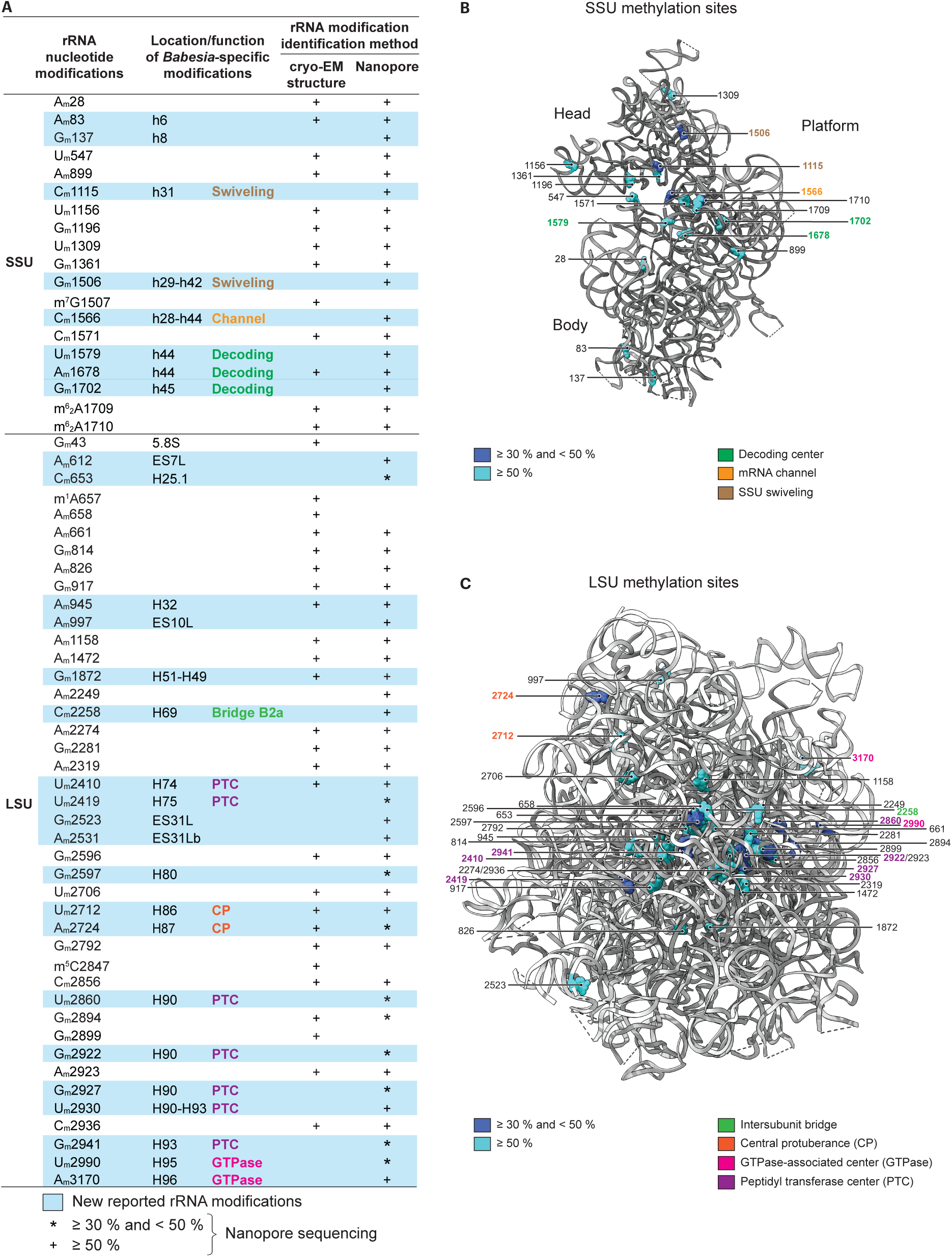
*Babesia* ribosome rRNA methylations. **A** rRNA modification list identified in this study, either by density map analysis (cryo-EM structure) and/or nanopore sequencing methods. For nanopore sequencing, methylations identified following an applied threshold of 50 % or higher are indicated by a plus sign (+). In comparison, those indicated with an asterisk (*) correspond to modifications identified following an applied threshold between 30 and 50 %. A hyphen (–) relates to a modification located between two rRNA helices (e.g., h29–h42). Nucleotides 612 and 2531 are unmodeled within the ribosome. **B** Methylation sphere representation in the 3D structure of the *Babesia* SSU, with a threshold set either between 30 and 50 %, or higher than 50 %. Base methylations and ribosomal proteins are omitted for clarity. **C** Methylation sphere representation in the 3D structure of the *Babesia* LSU, with a threshold set either between 30 and 50 %, or higher than 50 %. Ribosomal proteins and the 5.8S rRNA are omitted for clarity.

We observed methylations in the central protuberance (H86 and H87), in the intersubunit bridge B2a, located between the A- and P-tRNA binding sites (H69), and in the GTPase-associated center (H95 and H96) (**Figs. 5a and 5c**). We also detected a subset of rRNA methylations in the *Babesia* ESs (ES7L, ES10L, and ES31L, **Figs. 4 and 5a**).

Pseudouridylations are more challenging to identify in cryo-EM maps because they require high local resolution to observe their characteristic hydrogen bonding (**Fig. 2e**). Among the various methods available to identify pseudouridine modification positions^64,67^, we again turned to nanopore sequencing, as described above. We identified a total of 60 pseudouridylation sites, among which more than half were present at a threshold higher than 50 % (**Fig. 6**). The new pseudouridines in the SSU are generally located within the head, including in the mRNA channel, with a large group of pseudouridines found in the truncated ESs (ES3S, ES6S, ES7S, and in ES9S) (**Figs. 6a and 6b**). In the LSU, pseudouridines are generally clustered at the PTC, including in H92, which interacts with the CCA extremity of the A-site tRNA. Pseudouridines found in the line of the PTC (H90-H91) extend toward the surface, positioning them at the interface between the catalytic core and the binding site for translation factors. Modifications were also identified in inter-subunit bridges, including in H38 (A-site finger; bridge B1a) and in H69 (bridge B2a with SSU h44). Pseudouridines were also present in the truncated LSU ESs (ES7L, ES9L, and ES31L) (**Figs. 6a and 6c**). Thus, nearly half of the newly described modifications (methylations and pseudouridylations) are in functional centers, rather than in purely structural elements.

**Fig. 6.**
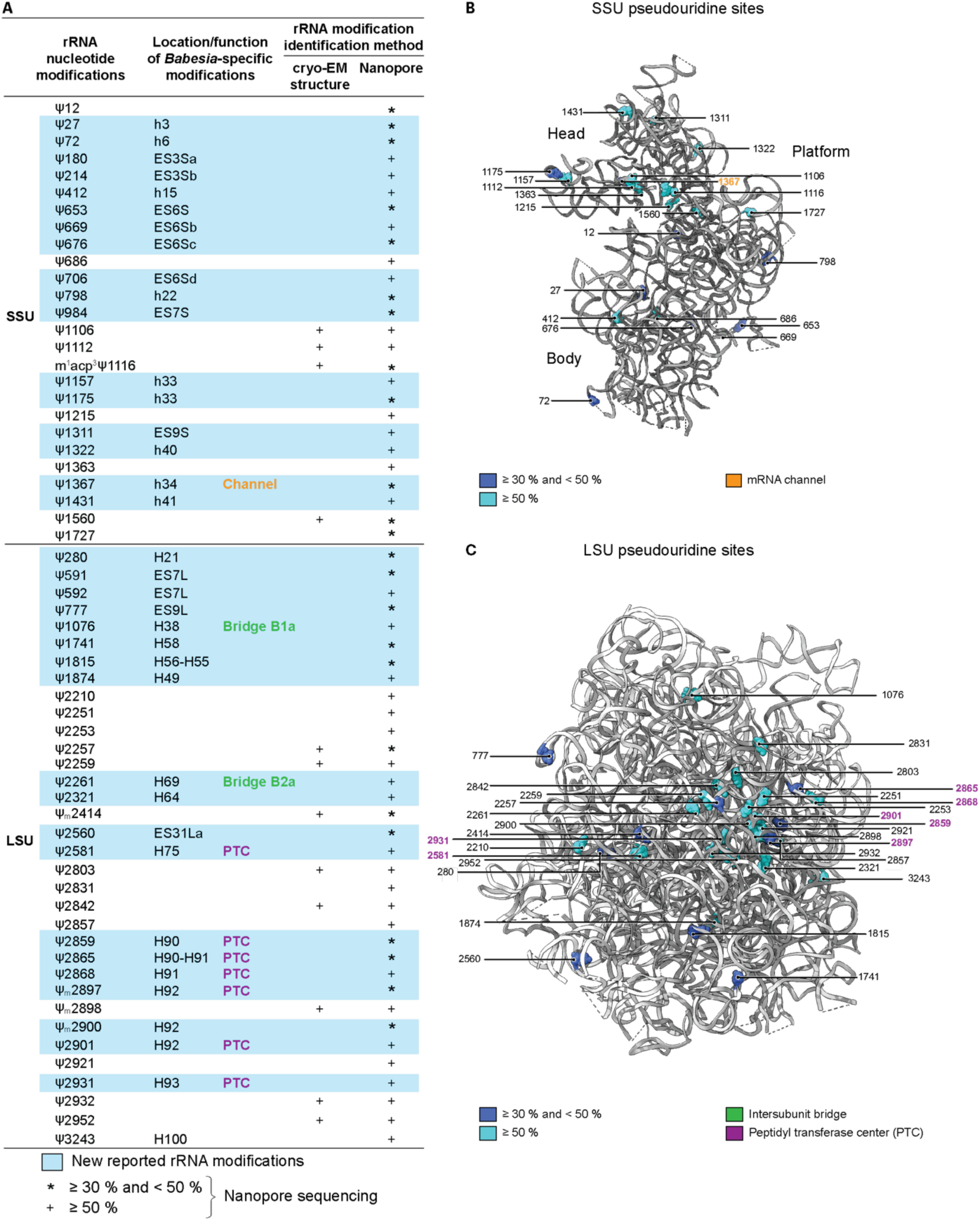
*Babesia* ribosome rRNA pseudouridylations. **A** rRNA modification list identified in this study, either by density map analysis (cryo-EM structure) and/or nanopore sequencing methods. For nanopore sequencing, pseudouridylations identified following an applied threshold of 50 % or higher are indicated by a plus sign (+). In com­ parison, those indicated with an asterisk (*) correspond to modifications identified following an applied threshold between 30 and 50 %. A hyphen(-) relates to a modification located between two helices (e.g., H90-H91). Nucleo­ tides 180,214,706,984, 591 and 592 are unmodeled within the ribosome. **B** Pseudouridylations sphere represen­ tation in the 3D structure of the *Babesia* SSU, with a threshold set either between 30 and 50 %, or higher than 50 %. Ribosomal proteins are omitted for clarity. **C** Pseudouridylations sphere representation in the 3D structure of the *Babesia* LSU, with a threshold set either between 30 and 50 %, or higher than 50 %. Ribosomal proteins and the 5.8S rRNA are omitted for clarity.

## Discussion

In this study, we present high-resolution cryo-EM structures of the *Babesia divergens* ribosome, and its landscape of rRNA modifications. Through our analyses of the two conformational states of the 80S ribosome, we show that the compact architecture of the *Babesia* rRNA, particularly within the LSU, results in significant remodeling of its surface, a change that may provide species-specific opportunities for translation regulation in *Babesia*. Along with the rRNA modification annotation for the *Babesia* ribosome, we report new modification sites enriched at functional and critical ribosome regions, such as the SSU decoding center and the LSU peptidyl transferase center.

High-resolution cryo-EM density maps of *B. divergens* ribosome enabled us to model all ribosomal proteins except those associated within the L1 and P stalks of the LSU^45–48^. Owing to the high-quality maps, we identified, in few instances, discrepancies between the density and the corresponding protein sequence from the *B. divergens* 1802 reference genome annotation. Consequently, we were able to correct these genome annotations in the VEuPathDB, which included a missing intron (e.g., uS7, eS10, uS12, eL18, and eL24), and the identification of previously unannotated genes (e.g., eS17, eL41). Moreover, the cryo-EM dataset showed no corresponding density for the ribosomal protein eL28, whereas this protein is present in *P. falciparum*^29,30,32^ and *T. gondii* cryo-EM ribosome structures^34^. In *P. falciparum*, eL28 interacts at the junction of two expansion segments, ES7La and ES15L, both of which are much shorter in *B. divergens*. We hypothesize that the reduction of the ESs in *B. divergens* may have led to the loss of eL28. The genomes of *B. microti* and *B. duncani* include a gene encoding eL28. Although the 28S rRNA is not yet annotated in *B. duncani*, we found that in *B. microti*, the ES7La rRNA is about the same length as in *P. falciparum*: this observation suggests that the ES7La contains the necessary rRNA elements to anchor eL28 on the *B. microti* ribosome. *B. divergens* has therefore simplified its ribosomal architecture over a short evolutionary period compared to *B. microti* and potentially *B. duncani*. These features collectively distinguish the *B. divergens* ribosome from those of other apicomplexans that may influence ribosomal assembly, stability, and translational control in this lineage.

Our insights into the composition and evolution of *B. divergens* ribosomal proteins prompted us to examine the association of other ribosome-bound factors that have previously shown divergent profiles among apicomplexan parasites. For example, we observed RACK1 associated with the head of the SSU, as previously described in yeast and human ribosomes^49,50^. This finding contrasts with previous studies in *Plasmodium* and *Toxoplasma*, in which density for RACK1 was absent or weak and unmodeled^29,30,33,34^. The robust presence of RACK1 may reflect either improved sample preservation during freezing and imaging, or the heterogeneity of the host-parasite life stages sampled, given that the parasites in this study were grown asynchronously. The latter suggestion is supported by a recent study in which *in situ* structures of ribosomes from cryo-FIB milled *P. falciparum*-infected human erythrocytes across all asexual stages show the presence of RACK1 on the SSU, although the lower resolution limited the detailed characterization of the interaction^32^. In contrast, a recent cryo-EM ribosome structure in *Plasmodium* suggests that artemisinin treatment increases the association of RACK1 with the SSU. Given the well-established biological importance of RACK1 association with ribosomes in other eukaryotic systems^68^, additional work will be required to delineate how RACK1 association with apicomplexan ribosomes influences their functional specialization.

In this study, we also present a comprehensive mapping of rRNA modifications in *Babesia*. By integrating nanopore sequencing with high-resolution cryo-EM structure determination, we identified an extensive set of methylation and pseudouridylation sites, including many that were previously unreported. We identified rRNA modifications at functional sites within the SSU, including the decoding center, mRNA channel, and intersubunit bridges b1a and b2a, as well as within the LSU, including the peptidyl transferase center and the GTPase-associated region. Several newly identified modifications are within the truncated ESs. Although the biological consequences of these modifications remain to be fully elucidated, their presence at functional hotspots points toward roles in modulating translation. Consistent with this observation, a single *Leishmania* pseudouridylation in H69 (Ψ512) (*L. donovani* and *L. major*) has been shown to be preferentially hypermodified in the host stage (amastigotes). This pseudouridylation affects tRNA selection, thereby increasing the translation of mRNAs that carry high-biased codons^24^. Interestingly, the corresponding site in *B. divergens* (Ψ2259) is also highly modified in our ribosome structure from parasites maintained in human blood culture, but whether it is hypomodified in the vector stage is unknown.

Beyond their potential impact on ribosome function and translation, unique rRNA modifications may also influence the susceptibility of *Babesia* ribosomes to small-molecule inhibitors. Several compounds targeting apicomplexan translation machinery have been described to date^69^. For example, mefloquine has been shown to bind the GTPase-associated center of the *Plasmodium* cytosolic ribosome, and certain derivatives exhibit enhanced potency by exploiting *P. falciparum*-specific ribosomal residues^35^. Emetine, a well-characterized translation inhibitor, binds to the ribosomal E-site and disrupts translocation, but its clinical application is limited by its high toxicity^30^. Similarly, pactamycin, another E-site binder, has inspired the development of analogs with reduced cytotoxicity that retain potent antimalarial activity^70^. Collectively, these studies highlight that the translational machinery is a promising yet underexplored target for antiparasitic drug development, analogous to strategies employed against bacterial pathogens. The divergent rRNA modifications observed in *Babesia* species relative to their human hosts may provide a unique opportunity for selective therapeutic targeting.

## Methods

### RIBOSOME SAMPLE PREPARATION

#### Parasite culture

We cultivated the *Babesia divergens* Rouen 1987 strain at high parasitemia (> 20 %), 37 °C, and 4 % hematocrit (HCT) in purified Caucasian male O-positive human red blood cells (RBCs from Research Blood Components) under low-oxygen conditions (1 % O_2_, 5 % CO_2_). We maintained cells in RPMI-1640 medium supplemented with 25 mM HEPES, 367.5 µM hypoxanthine, and 25 µg/mL gentamicin at pH 6.75. This medium was then supplemented with 22.2 mM sodium bicarbonate and 4.31 mg/mL AlbuMAX II (Invitrogen). We isolated parasites from RBCs by mechanical release, followed by filtration through a 1.2 µm Luer-lock syringe filter and centrifugation at 3000 x g for five minutes^71,72^. We processed a 30 mL culture volume (4 % HCT, 1.2 mL packed RBCs) according to this procedure, which we repeated for a total of three liters of culture volume. To isolate free merozoites contained in the supernatant, we centrifuged at 4100 xg for 15 minutes.

#### Ribosome purification

We washed parasite pellets twice with 1X PBS at 16,000 x g for 5 minutes. We then resuspended the pellet in a lysis buffer (LB: 20 mM HEPES pH 7.4, 300 mM KCl, 25 mM Mg(CH_3_COO)_2_, 0.15 % Triton X-100, 5 mM 2-β-mercaptoethanol, and complete protease inhibitor tablet from Roche), incubated at 4 °C for one hour, followed by two freeze-thaw cycles. We centrifuged the lysates at 12,000 x g for 10 minutes. We loaded the supernatant on a sucrose cushion buffer (SCB: 20 mM HEPES pH 7.4, 150 mM KCH_3_COO, 10 mM NH_4_CH_3_COO, 10 mM Mg(CH_3_COO)_2_, 30 % (w/v) sucrose, and 5 mM 2-β-mercaptoethanol). We centrifuged at 4 °C for 16 hours at 40,000 rpm in a 70.1 Ti rotor (Beckman Coulter). We resuspended the crude ribosomal pellet in 300 µL of ribosome buffer (RB: SCB devoid of sucrose) and then centrifuged it at 14,000 x g for 15 minutes. We layered the supernatant onto a 10-40 % (w/v) sucrose gradient (Biocomp Gradient Master) and centrifuged at 4°C for 3 hours at 41,000 rpm using an SW41 Ti rotor (Beckman Coulter). We collected and pooled the fractions containing ribosomes according to the UV absorbance profile (Biocomp Gradient Fractionator equipped with a Triax flow cell). We back-exchanged the buffer into the RB described above using an Amicon Ultra-4 centrifugal filter unit with a 50 kDa cutoff (Millipore).

### CRYO-EM DATA COLLECTION AND PROCESSING

#### Data collection by cryo-electron microscopy

We glow-discharged cryo-EM grids (Quantifoil R1.2/1.3 with Ultrathin Carbon) using a Pelco Easiglow plasma discharge system (Ted Pella) at 15 mA for 30 seconds. We plunge-freeze glow-discharged cryo-EM grids, previously incubated with 3 µL of *B. divergens* ribosomes (∼100 nM) for 1 minute, using a FEI Vitrobot Mark IV (Thermo Fisher Scientific) operating at 100 % humidity, 4 °C, and a blot force of 12 for 3 sec. We then transferred the grids to an FEI Titan Krios electron microscope, operated at 300 kV, using Serial EM for automated data acquisition^73^. We collected one data set (**Tables S1, S2**) at a nominal underfocus of −0.8 to −2.2 µm at a magnification of 105,000 x, yielding a pixel size of 0.825 Å. We recorded the micrographs as a movie stack (12,114 frame stacks/movies) on a K3 direct electron detector (Gatan) operated in counting mode. We collected 40 frames per 1.4-second movie, with a total dose of 42 e/Å^2^.

#### Single-particle image processing (Figs. S1, S2)

We gain corrected, aligned (5 x 5 patch), and dose-weighted the dose-fractionated movies using MotionCor2^74^. We used the unweighted, aligned micrographs for contrast transfer function (CTF) estimation with CTFFIND-4.1.13^75^. We used crYOLO to select 1,259,788 particles^76^. We used RELION 3.1.2 for 2D and 3D classification and refinements^77^. We extracted the particles with a box size of 448 pixels (370 Å) and binned them fourfold. We split the particles into three subsets (size split 500000), and each subset was subjected to two rounds of 2D classification. We used a subset of the well-resolved 2D class averages (418,357 particles) for the ab initio 3D reconstruction model. We used this model as a starting reference (filtered to 40 Å) for a 3D consensus refinement. We then subjected each subset to 3D classification without alignments and selected 3D classes that showed well-formed ribosome particles. We created a mask of the inter-subunit space (PDB 6GZ3)^78^ and carried out focused classification. We reextracted the particles at the original pixel size. We refined each class independently using an initial angular sampling set of 15 degrees and local searches from an auto sampling set at 3.7 degrees. Three classes displayed clear morphology for 80S particles. We initially selected the most stable and populated class (class 3) and did per-particle CTF refinement, aberration refinement, followed by signal-subtracted focused refinement and classification without alignments. We then did another round of 3D auto-refinement. Before sharpening (post-processing), we created a mask for the 80S using a lowpass filter of 15 Å and an initial binarization threshold of 0.003. We extended the binary map by 4 pixels in every direction and added a cosine-shaped soft-edge of 4 pixels, resulting in an 80S map with a global resolution of 2.6 Å. We performed a multibody refinement to improve the resolution of the distinct subdomains, focusing on the SSU head and body and the LSU regions separately. We combined the resulting focus-refined maps using ChimeraX^37^ to generate a full 80S map. We also further processed a class (class 1) containing density for E-, P-, and A-site tRNAs and mRNA. Briefly after a round of 3D auto-refinement, we performed a focused classification (without alignments) of the tRNAs to isolate a more homogeneous subset, followed by another round of 3D auto-refinement. We then performed CTF and aberration refinement, and re-refined and post-processed the map using an 80S mask (lowpass filtered at 15 Å, binarization threshold 0.004, 4-pixel extension, and 4-pixel soft edge), yielding a global resolution of 3.1 Å. Multibody refinement was also used to improve the resolution of individual subdomains, and a composite map was generated for this class.

#### Atomic model building and refinement

We retrieved ribosomal protein sequences from *B. divergens* 1802A (the reference strain) using *P. falciparum* 3D7 sequences (PDB IDs: 3J7A, 3J79)^30^ and the BLAST tool from PiroplasmaDB and PlasmoDB, a part of the VEuPathDB^79^. We input the extracted *B. divergens* sequences into Alphafold2 using the Colab server with template and amber relaxation options enabled^36^. We used AlphaFold2-validated metrics to assess model reliability and selected the best-relaxed structures with the highest pLDDT for docking into the post-processed *B. divergens* maps using ChimeraX^37^. We retrieved rRNA sequences from *B. divergens* using the RNAcentral database^38^. We aligned the sequences with those of *T. gondii* structures (PDB IDs: 5XXB, 5XXU)^34^ using Clustal Omega^39^. We used predicted secondary structure maps from RNAcentral to aid in rRNA modeling and mutated individual rRNA residues based on the *B. divergens* sequence. We built *de novo* less conserved regions and linker sequences using Coot^80^. We reviewed base-pairing after manual adjustment with Coot. We initially focused on building the atomic model corresponding to ribosomes bound to an E-site tRNA (conformational state 2), given the highest-resolution maps. We refined the atomic model against the corresponding density map using real-space refinement (phenix.real_ space_refine) in PHENIX^41^. We applied iterative rounds of manual model adjustment in Coot and model refinement in PHENIX. The E-site tRNA (PDB 3J7A)^30^ was docked into the ribosome using ChimeraX and further refined in PHENIX. We evaluated the quality of the final model using MolProbity^81^. We docked this model into 3D class 1, docked the P, and A-site tRNAs (PDB 9BUP) followed by manual rebuilding of the rRNA. We used ModelAngelo^82^ for generating ribosomal protein coordinates that showed local rearrangements when compared to the 3D class 3. The model was further refined with iterative rounds of manual adjustment in Coot and real-space refinement as described above. Based on our atomic ribosome models, we improved the VEuPathDB annotation set in PiroplasmaDB through manual curation in Apollo^79^, by correcting misannotations of gene models coding for ribosomal proteins uS7, uS10, uS12, EL15, EL18, eL24, and eL41.

### MASS SPECTROMETRY ANALYSIS

#### Sample preparation

We denatured ribosomal proteins with Laemmli buffer supplemented with 2-β-mercaptoethanol to *B. divergens* ribosomes (1:1 ratio), previously purified, and boiled at 100 °C for three minutes. We precipitated the proteins by adding five volumes of 0.1 M ammonium acetate in 100 % methanol to the Laemmli-containing samples. We centrifuged the samples at 12,000 x g at 4 °C for 15 min. We washed the resulting protein pellet twice with 0.1 M ammonium acetate in 80 % methanol and dried the pellet under vacuum (Speed-Vac concentrator). We submitted the dried pellet to the Harvard Center for Mass Spectrometry (HCMS). A dried protein sample of 5 µg was dissolved in 100 µL of 50 mM triethylammonium bicarbonate (TEAB) and reduced by the addition of 20 mM tris(2-carboxyethyl) phosphine (TCEP) and incubated at 37 °C for 45 minutes. Proteins were alkylated in 40 mM iodoacetamide for 45 min in the dark at room temperature. Proteins were trypsinized at a 1:50 ratio (w/w, substrate/enzyme) and incubated overnight at 37 °C. Digested samples were cleaned with C18 Spin Tips (Pierce, Thermo Scientific) and then reconstituted with 0.1 % formic acid before LC-MS/MS analysis.

#### LC-MS/MS analysis

LC-MS/MS experiments were performed on a Lumos Tribrid Orbitrap Mass Spectrometer equipped with Ultimate 3000 (Thermo Fisher) nano-HPLC. Peptides were separated on a 150 μm inner diameter microcapillary trapping column packed with approximately 2 cm of C18 Reprosil resin (5 μm, 100 Å, Dr. Maisch GmbH, Germany), followed by an analytical column (µPAC, C18 pillar surface, 50 cm bed, Thermo Scientific). Separation was achieved by applying a 5–35 % acetonitrile gradient in 0.1 % formic acid over 180 minutes at a flow rate of 300 nL/min. Electrospray ionization occurred with a voltage of 2 kV using a homemade electrode junction at the end of the microcapillary column, sprayed from metal tips (PepSep, Denmark). The mass spectrometry survey scan was performed in the Orbitrap in the range of 410–1,800 m/z at a resolution of 12×10^4^, followed by the selection of the twenty most intense ions (TOP20) for CID-MS2 fragmentation in the Ion trap using a precursor isolation width window of 2 m/z, AGC setting of 10,000, and a maximum ion accumulation of 100 ms. Singly charged ion species were not subjected to CID fragmentation. The normalized collision energy was set to 35 V with an activation time of 10 ms. Ions in a ten-ppm m/z window around ions selected for MS2 were excluded from further selection for fragmentation for 60 seconds. Raw data from LC-MS/MS experiments were submitted for analysis in Proteome Discoverer 2.4 software (Thermo Scientific). The MS/MS data were searched against the customized protein sequence (*B. divergens* dataset from PiroplasmaDB) and against known contaminants, including human keratins and typical laboratory contaminants. Sequest HT searches were performed using the following guidelines: a 15 ppm MS tolerance and 0.6 Da MS/MS tolerance; trypsin digestion with up to two missed cleavages; carbamidomethyl group on cysteine as static modification; oxidation of methionine, N-terminal acetylation, and deamidation of asparagine and glutamine as variable modifications; and required peptide length ≥ 6 amino acids. At least one unique peptide per protein group is needed to identify proteins. An MS2 spectra assignment FDR of 1 % on both protein and peptide levels was achieved by applying a target-decoy database search with Percolator.

### RNA ANALYSIS

#### RNA isolation and sequencing

We cultured and harvested the parasites as described above. A 50 mL culture volume of *B. divergens* at 2 % HCT and ∼ 15-20 % parasitemia. The RBCs were resuspended in 0.33 % saponin to release parasites, and free merozoites were also collected from the supernatant by additional rounds of centrifugation. Pellets were washed in Dulbecco’s PBS (DPBS) and homogenized in TRI Reagent (ThermoFisher). Total RNA was extracted from *B. divergens* using a Direct-zol RNA Miniprep Kit (Zymo Research) according to the manufacturer’s instructions. We performed on-column DNase I treatment for 20 minutes at room temperature. We assessed the concentration and quality of the total RNA using a NanoDrop spectrophotometer and RNA Flash Gel Fragment Separation (Lonza).

#### Oxford Nanopore sequencing

We targeted *B. divergens* 28S and 18S rRNAs using custom oligos complementary to their specific 3′ ends (18S: 5’-GAGGCGAGCGGTCAATTTTCCTAAGAGCAAGAAGAAGCCGAATGATCCT-3’, 28S: 5’-GAGGCGAGCGGTCAATTTTCCTAAGAGCAAGAAGAAGCCATACACACAA-3’). These target-specific oligos were annealed to an adapter oligo (5’-/5PHOS/GGCTTCTTCTTGCTCTTAGGTAGTAGGTTC-3’) to generate functional reverse transcriptase adaptors (RTAs) according to Oxford Nanopore Technologies (ONT) sequence-specific SQK-RNA004 protocol. Briefly, the adaptor oligo and target-specific oligo were mixed in a 1:1 molar ratio at a final concentration of 1.4 µM in an annealing buffer (10 mM Tris-HCl, 50 mM NaCl, pH 7.5), heated to 95 °C for 2 minutes, and allowed to cool slowly to room temperature. We submitted the extracted total RNA samples described above, along with the custom RTAs, to the Harvard University Bauer Core Facility. The facility evaluated the precise concentration, RNA integrity, and quality using QuBit and TapeStation before proceeding to RNA library preparation (SQK-RNA004 kit) and nanopore sequencing. The 28S or 18S RNA libraries were loaded onto a PromethION flow cell (FLO-PRO004RA) twice (2 runs). The raw signals were recorded and stored in POD5 format. Real-time basecalling was enabled in MinKNOW v24.06.15, using the high-accuracy model (v3.0.1). Reads were filtered with a minimum quality (Q) score ≥ 9.

#### RNA modification analysis and statistics

We rebasecalled the raw sequencing signals (POD5) using Oxford Nanopore’s basecaller Dorado 0.9.0 in super-accurate mode (rna004_130pbs_sup@v5.1.0) with pseudorudine base detection enabled. We used Dorado v1.0.2 (*rna004_130bps_sup@v5.2.0*) to additionally detect 2′-O-methylations, a feature newly introduced in this version of the basecaller. We mapped the base-called reads to the *B. divergens* 18S or 28S rRNA using the Dorado aligner. The aligned reads were filtered, sorted, and indexed using SAMtools^83^. We used Modkit (v0.5.0) to identify modification sites, aggregate per-read calls, and compute counts of modified (Nmod) and unmodified reads at each position. Modkit automatically estimated a confidence threshold from a subset of the reads (10th percentile of the site-specific probability distribution) to exclude low-confidence basecalls, with per-site coverage capped (maximum depth) at 7.5 million reads. In addition to the standard *% Modified* reported by Modkit (Nmod/Nvalid_cov × 100), we calculated an adjusted modification percentage defined as Nmod / (Nvalid_cov + Ndiff) × 100. The Ndiff counts, provided directly by Modkit, represent reads at that position that do not contain the nucleotide evaluated by the model and are therefore excluded from the default calculation (Nvalid_cov), even when such reads make up the majority of coverage. Including Ndiff in the denominator prevents inflated values at sites where the nucleotide subject to modification is rare.

## Data availability

Atomic coordinates and cryo-EM maps have been deposited in the PDB and Electron Microscopy Data Bank (EMDB), respectively. For the conformational state 1 (3DC1, 80S ribosome with three tRNAs and mRNA), the PDB entry is 9YXB with the following associated EMDB entries: EMD-73602 (composite), EMD-73574 (consensus), EMD-73576 (LSU), EMD-73577 (SSU body), EMD-73581 (SSU head). For the conformational state 2 (3DC3, 80S ribosome with E-site tRNA), the PDB entry is 9YGM (3DC3, 80S ribosome with E-site tRNA) with the following associated EMDB entries: EMD-72933 (composite), EMD-72052 (consensus), EMD-72055 (LSU), EMD-72069 (SSU body), EMD-72070 (SSU head). The mass spectrometry proteomics data have been deposited to the ProteomeXchange Consortium via the PRIDE partner repository with the dataset identifier PXD069329 and 10.6019/PXD069329. Nanopore sequencing datasets have been deposited in NCBI SA with accession IDS: PRJNA1358264 and SAMN53107339. All other materials are available upon request and should be addressed to MLA.

## Acknowledgements

We thank Drs. K. Deitsch and L. Kirkman (Weill Cornell Medical College) for providing the *B. divergens* Rouen 1987 strain. The authors thank the Molecular Electron Microscopy Suite at Harvard Medical School, particularly Dr. S. Sterling, for her assistance in cryo-EM data collection. We thank Dr. B. Budnik and R. Robinson at the Harvard Center for Mass Spectrometry for their assistance with the proteomics analyses. The authors thank C. Hartman and K. Cribari from the Bauer Core Facility at Harvard for library preparation and Oxford nanopore sequencing. The author thanks Dr. O. Harb from VEuPathDB’s team for his assistance with genome sequence analysis. We also thank Drs. J. Abraham and S.C. Harrison from Harvard Medical School for their scientific discussion and critical reading of the manuscript.

## Author contributions

Conceptualization: MLA

Methodology: MLA, MTD, CGV, LSIT, CDK

Investigation: CGV, CDK, LSIT, DCC, PC

Supervision: MLA, MTD

Provided reagents: MLA, MTD

Formal analysis: MLA, MTD, CGV, LSIT

Funding acquisition: MLA, MTD

Writing – original draft: MLA, CGV

Writing – final draft: All authors

