## Supplementary Tables and Figures for "Ribosomal Architecture and rRNA Modification Landscape in the Tick-Borne Parasite *Babesia divergens*"

**Table S1 Cryo-EM data collection, refinement, and validation statistics for ribosomes with an A-, P-, E-site tRNAs and an mRNA fragment (conformational state 1, related to Figs. 1 and 2).**

| | | | | | |
|---|---|---|---|---|---|
| **Data collection and processing** | | | | | |
| Microscope | FEI Titan Krios | | | | |
| Detector | K3 Summit Direct Electron Detector | | | | |
| Detector mode | Counting | | | | |
| Voltage (kV) | 300 | | | | |
| Magnification | 105,000 x | | | | |
| Pixel size (Å) | 0.825 | | | | |
| Defocus range (μm) | -0.8 to -2.2 | | | | |
| Electron exposure (e-/Å$^2$) | 42 | | | | |
| Number of frames per movie | 40 | | | | |
| Exposure rate (e-/Å$^2$/frame$^{-1}$) | 1.05 | | | | |
| Data collection software | SerialEM 3.8.6 | | | | |
| Data processing software | RELION (versions 3.1.3 and 5.0) | | | | |
| Usable micrographs | 12, 097 | | | | |
| Particles extracted | 1,259,788 | | | | |
| Particles used for reconstruction | 26,467 | | | | |
| FSC threshold | 0.143 | | | | |
| EMDB accession code (EMD) | 73574 | 73576 | 73577 | 73581 | 73602 |
| Map Description | Consensus | LSU | SSU body | SSU head | Composite |
| Map resolution (Å) global | 3.1 | 3.0 | 3.0 | 3.1 | - |
| Unmasked ( Å, 0.143 criterion) | 3.6 | 3.5 | 3.9 | 4.0 | - |
| Masked ( Å, 0.143 criterion ) | 3.1 | 3.0 | 3.0 | 3.1 | - |
| PDB accession code (80S) | 9YXB | | | | |
| **Refinement Statistics** | | | | | |
| Initial model(s) used | PDB 9YGM (conformation 2 with an E-site tRNA) | | | | |
| Refinement package | PHENIX 1.21.1-5286 | | | | |
| Model resolution (Å, 0.5 criterion) | 3.2 | | | | |
| Map sharpening *B* factor (Å) | LSU (75.1), SSU body(74.0), SSU head (76.1) | | | | |
| Map CC | 0.81 | | | | |
| Model composition | | | | | |
| Number of chains | 82 | | | | |
| Non-hydrogen atoms | 186900 | | | | |
| Residues | 10229 | | | | |
| Nucleotides | 4913 | | | | |
| Ligands/ions | K:21, MG:135 | | | | |
| Water | 0 | | | | |
| *B* factors (Å$^2$) (min/max/mean) | | | | | |
| Proteins | 26.82/221.86/78.71 | | | | |
| Nucleotides | 0.00/768.38/85.49 | | | | |
| R.m.s. deviations | | | | | |
| Bond lengths (Å) | 0.005 | | | | |
| Bond angles (°) | 0.774 | | | | |
| **Validation Statistics** | | | | | |
| MolProbity score | 2.07 | | | | |
| Clash score | 8 | | | | |
| Rotamer outliers (%) | 3 | | | | |
| C-beta deviations | 0 | | | | |
| CaBLAM outliers | 1.78 | | | | |
| Ramachandran plot | | | | | |
| Favored (%) | 95.93 | | | | |
| Allowed (%) | 3.97 | | | | |
| Outliers (%) | 0.10 | | | | |

**Table S2 Cryo-EM data collection, refinement, and validation statistics for ribosomes with an E-site tRNA (conformational state 2, related to Figs. 1 and 2).**

| | | | | | |
|---|---|---|---|---|---|
| **Data collection and processing** | | | | | |
| Microscope | FEI Titan Krios | | | | |
| Detector | K3 Summit Direct Electron Detector | | | | |
| Detector mode | Counting | | | | |
| Voltage (kV) | 300 | | | | |
| Magnification | 105,000 x | | | | |
| Pixel size (Å) | 0.825 | | | | |
| Defocus range (μm) | -0.8 to -2.2 | | | | |
| Electron exposure (e-/Å$^2$) | 42 | | | | |
| Number of frames per movie | 40 | | | | |
| Exposure rate (e-/Å$^2$/frame$^{-1}$) | 1.05 | | | | |
| Data collection software | SerialEM 3.8.6 | | | | |
| Data processing software | RELION 3.1.3 | | | | |
| Usable micrographs | 12, 097 | | | | |
| Particles extracted | 1,259,788 | | | | |
| Particles used for reconstruction | 261,009 | | | | |
| FSC threshold | 0.143 | | | | |
| EMDB accession code (EMD) | 72052 | 72055 | 72069 | 72070 | 72933 |
| Map Description | Consensus | LSU | SSU body | SSU head | Composite |
| Map resolution (Å) global | 2.6 | 2.5 | 2.6 | 2.7 | - |
| Unmasked (Å, 0.143 criterion) | 2.8 | 2.8 | 2.9 | 3.1 | - |
| Masked (Å, 0.143 criterion) | 2.6 | 2.5 | 2.6 | 2.7 | - |
| PDB accession code (80S) | 9YGM | | | | |
| **Refinement Statistics** | | | | | |
| Initial model(s) used | Alphafold2 (ribosomal proteins), PDB 5XXB and 5XXU (rRNA) | | | | |
| Refinement package | PHENIX 1.21.1-5286 | | | | |
| Model resolution (Å, 0.5 criterion) | 2.74 | | | | |
| Map sharpening *B* factor (Å) | LSU (93.9), SSU body(60.1), SSU head (64.5) | | | | |
| Map CC | 0.86 | | | | |
| Model composition | | | | | |
| Number of chains | 79 | | | | |
| Non-hydrogen atoms | 187170 | | | | |
| Residues | 10664 | | | | |
| Nucleotides | 4762 | | | | |
| Ligands/ions | ZN: 5, K: 25, MG: 136, IAS:1 | | | | |
| Water | 22 | | | | |
| *B* factors (Å$^2$) (min/max/mean) | | | | | |
| Proteins | 24.56/284.38/73.23 | | | | |
| Nucleotides | 4.41/407.80/68.19 | | | | |
| R.m.s. deviations | | | | | |
| Bond lengths (Å) | 0.007 | | | | |
| Bond angles (°) | 0.648 | | | | |
| **Validation Statistics** | | | | | |
| MolProbity score | 1.66 | | | | |
| Clash score | 6 | | | | |
| Rotamer outliers (%) | 3 | | | | |
| C-beta deviations | 0 | | | | |
| CaBLAM outliers | 1.40 | | | | |
| Ramachandran plot | | | | | |
| Favored (%) | 97.77 | | | | |
| Allowed (%) | 2.21 | | | | |
| Outliers (%) | 0.02 | | | | |

**Table S3 Reference table of the *Babesia* ribosomal proteins from the small subunit (related to Fig. 1)**. The chain IDs correspond to the ribosomal proteins modeled in the 80S ribosome structure as identified by the PDB code 9YGM. The sequence alignment identity percentage is provided for comparison between *B. divergens* 1802A and *P. falciparum* 3D7, or between *B. bovis* T2Bo and PF3D7, for eS17.

| Gene ID (*Babesia*) | Unified r-prot. name | Chain ID | Number of residues modeled/sequence | Gene ID (*Plasmodium*) | Sequence residue number | Sequence alignment identity (%) |
|---|---|---|---|---|---|---|
| Bdiv_040040 | eS1 | SA | 214/264 | PF3D7_0322900 | 262 | 66 |
| Bdiv_033120 | uS2 | SB | 196/274 | PF3D7_1026800 | 263 | 67 |
| Bdiv_011570 | uS3 | SC | 209/223 | PF3D7_1465900 | 221 | 79 |
| Bdiv_010520c | uS4 | SD | 178/184 | PF3D7_0520000 | 189 | 85 |
| Bdiv_034010 | eS4 | SE | 260/266 | PF3D7_1105400 | 261 | 60 |
| Bdiv_029510 | uS5 | SF | 180/196 | PF3D7_1447000 | 272 | 80 |
| Bdiv_037560c | eS6 | SG | 216/239 | PF3D7_1342000 | 306 | 62 |
| Bdiv_017820c | uS7 | SH | 178/220 | PF3D7_0721600 | 195 | 74 |
| Bdiv_032090 | eS7 | SI | 178/194 | PF3D7_1302800 | 194 | 51 |
| Bdiv_016440c | uS8 | SJ | 129/130 | PF3D7_0316800 | 130 | 79 |
| Bdiv_012600 | eS8 | SK | 181/192 | PF3D7_1408600 | 218 | 54 |
| Bdiv_027730 | uS9 | SL | 134/149 | PF3D7_0813900 | 144 | 62 |
| Bdiv_009240 | uS10 | SM | 95/120 | PF3D7_1003500 | 118 | 59 |
| Bdiv_032170 | eS10 | SN | 96/134 | PF3D7_0719700 | 137 | 54 |
| Bdiv_010180c | uS11 | SO | 125/157 | PF3D7_0516200 | 151 | 92 |
| Bdiv_040390c | uS12 | SP | 186/188 | PF3D7_0306900 | 145 | 66 |
| Bdiv_040260c | eS12 | SQ | 115/135 | PF3D7_0307100 | 141 | 47 |
| Bdiv_027650 | uS13 | SR | 144/154 | PF3D7_1126200 | 156 | 71 |
| Bdiv_036780 | uS14 | SS | 50/66 | PF3D7_0705700 | 54 | 65 |
| Bdiv_001750c | uS15 | ST | 149/151 | PF3D7_1358800 | 151 | 67 |
| Bdiv_008760 | uS17 | SU | 151/156 | PF3D7_0317600 | 161 | 78 |
| BBOV_III001250* | eS17 | SV | 77/132 | PF3D7_1242700 | 137 | 73 |
| Bdiv_002790 | uS19 | SW | 118/149 | PF3D7_1317800 | 145 | 67 |
| Bdiv_040250 | eS19 | SX | 146/174 | PF3D7_0422400 | 170 | 47 |
| Bdiv_033240c | eS21 | SY | 78/79 | PF3D7_1144000 | 82 | 55 |
| Bdiv_010070 | eS24 | SZ | 96/135 | PF3D7_0519400 | 133 | 60 |
| Bdiv_022980c | eS25 | Sa | 72/104 | PF3D7_1421200 | 105 | 48 |
| Bdiv_038810c | eS26 | Sb | 97/115 | PF3D7_0217800 | 107 | 70 |
| Bdiv_036570 | eS27 | Sc | 70/82 | PF3D7_1308300 | 82 | 72 |
| Bdiv_019090c | eS28 | Sd | 63/67 | PF3D7_1461300 | 67 | 82 |
| Bdiv_039110c | eS30 | Se | 40/60 | PF3D7_0219200 | 58 | 72 |
| Bdiv_033080 | eS31 | Sf | 60/157 | PF3D7_1402500 | 149 | 46 |
| Bdiv_010290 | RACK1 | Sg | 322/323 | PF3D7_0826700 | 323 | 63 |

* *B. bovis* corresponding sequence in *B. divergens* Rouen 1987 is within CCSG02000001.1: 1890202..1890772 supercontig.

**Table S4 Reference table of the *Babesia* ribosomal proteins from the large subunit (related to Figs. 1 and 3)**. The chain IDs correspond to the ribosomal proteins modeled in the 80S ribosome structures as identified by the PDB code 9YGM. The sequence alignment identity percentage is provided for comparison between *B. divergens* 1802A and *P. falciparum* 3D7, or between *T. gondii* ME49 and PF3D7, for eL41.

| Gene ID (*Babesia*) | Unified r-prot. name | Chain ID | Number of residues modeled/sequence | Gene ID (*Plasmodium*) | Sequence residue number | Sequence alignment identity (%) |
|---|---|---|---|---|---|---|
| Bdiv_038620 | uL1 | N.M. | 0/217 | PF3D7_1441200 | 217 | 65 |
| Bdiv_015470 | uL2 | LA | 248/257 | PF3D7_0516900 | 260 | 69 |
| Bdiv_003320c | uL3 | LB | 381/395 | PF3D7_1027800 | 386 | 74 |
| Bdiv_004040c | uL4 | LC | 343/360 | PF3D7_0507100 | 411 | 50 |
| Bdiv_027640 | uL5 | LD | 166/171 | PF3D7_0719600 | 173 | 79 |
| Bdiv_027090 | uL6 | LE | 186/190 | PF3D7_1323100 | 190 | 64 |
| Bdiv_030770c | eL6 | LF | 149/212 | PF3D7_1338200 | 221 | 35 |
| Bdiv_016730 | eL8 | LG | 224/285 | PF3D7_1424400 | 283 | 53 |
| Bdiv_028610c | uL10 | N.M. | 0/313 | PF3D7_1130200 | 316 | 57 |
| Bdiv_010230c | uL11 | N.M. | 0/165 | PF3D7_0517000 | 165 | 66 |
| Bdiv_010650 | uL13 | LH | 201/202 | PF3D7_1004000 | 202 | 63 |
| Bdiv_027720c | eL13 | LI | 196/203 | PF3D7_0814000 | 215 | 43 |
| Bdiv_035030 | uL14 | LJ | 129/139 | PF3D7_1331800 | 139 | 80 |
| Bdiv_023300 | eL14 | LK | 128/133 | PF3D7_1431700 | 165 | 51 |
| Bdiv_034950 | uL15 | LL | 146/147 | PF3D7_0618300 | 148 | 64 |
| Bdiv_018330 | eL15 | LM | 198/198 | PF3D7_0415900 | 205 | 73 |
| Bdiv_016810c | uL16 | LN | 212/227 | PF3D7_1414300 | 219 | 72 |
| Bdiv_016720c | uL18 | LO | 289/306 | PF3D7_1424100 | 294 | 53 |
| Bdiv_037540 | eL18 | LP | 186/247 | PF3D7_1341200 | 184 | 53 |
| Bdiv_001430 | eL19 | LQ | 175/194 | PF3D7_0614500 | 182 | 63 |
| Bdiv_037530 | eL20 | LR | 184/188 | PF3D7_1341200 | 184 | 57 |
| Bdiv_016780c | eL21 | LS | 154/160 | PF3D7_1426000 | 161 | 46 |
| Bdiv_004980c | uL22 | LT | 152/194 | PF3D7_1351400 | 203 | 56 |
| Bdiv_011350c | eL22 | LU | 97/122 | PF3D7_0821700 | 139 | 58 |
| Bdiv_039940 | uL23 | LV | 118/153 | PF3D7_1323400 | 190 | 53 |
| Bdiv_032900 | uL24 | LW | 123/137 | PF3D7_0312800 | 126 | 61 |
| Bdiv_036590c | eL24 | LX | 61/199 | PF3D7_1309100 | 162 | 50 |
| Bdiv_016640 | eL27 | LY | 138/146 | PF3D7_1460700 | 146 | 50 |
| N.A. | eL28 | N.D. | - | PF3D7_1142500 | 127 | - |
| Bdiv_020740c | uL29 | La | 119/123 | PF3D7_1124900 | 124 | 46 |
| Bdiv_016690 | eL29 | Lb | 53/59 | PF3D7_1460300 | 67 | 70 |
| Bdiv_040380c | uL30 | Lc | 216/259 | PF3D7_0307200 | 257 | 55 |
| Bdiv_034830c | eL30 | Ld | 93/108 | PF3D7_1019400 | 108 | 61 |
| Bdiv_021180 | eL31 | Le | 105/117 | PF3D7_0503800 | 120 | 55 |
| Bdiv_038500 | eL32 | Lf | 125/132 | PF3D7_0903900 | 131 | 53 |
| Bdiv_039460 | eL33 | Lg | 108/115 | PF3D7_1142600 | 140 | 45 |
| Bdiv_019820c | eL34 | Lh | 100/150 | PF3D7_0710600 | 150 | 46 |

**Table S5 Ribosomal proteins identification by LC-MS/MS from the *Babesia* small subunit (related to Fig. 1)**. Sample purified from 80S ribosomes as identified by the PDB code 9YGM.

| **Gene ID *B. divergens* 1802A** | **Unified r-prot. name** | **Coverage (%)** | **Length (amino acids)** | **Molecular weight (kDa)** | **Number of peptides** | **Spectral counts** |
|---|---|---|---|---|---|---|
| Bdiv_040040 | eS1 | 49 | 264 | 30.0 | 15 | 216 |
| Bdiv_033120 | uS2 | 27 | 274 | 31.0 | 6 | 89 |
| Bdiv_011570 | uS3 | 53 | 223 | 24.7 | 10 | 188 |
| Bdiv_010520c | uS4 | 35 | 184 | 21.6 | 8 | 70 |
| Bdiv_034010 | eS4 | 31 | 266 | 30.1 | 8 | 114 |
| Bdiv_029510 | uS5 | 30 | 196 | 21.1 | 5 | 48 |
| Bdiv_037560c | eS6 | 44 | 239 | 27.5 | 11 | 168 |
| Bdiv_017820c | uS7 | 38 | 220 | 24.7 | 10 | 79 |
| Bdiv_032090 | eS7 | 18 | 194 | 22.0 | 8 | 69 |
| Bdiv_016440c | uS8 | 13 | 130 | 14.8 | 2 | 12 |
| Bdiv_012600 | eS8 | 54 | 192 | 21.9 | 9 | 121 |
| Bdiv_027730 | uS9 | 44 | 149 | 16.7 | 7 | 52 |
| Bdiv_009240 | uS10 | 36 | 120 | 13.5 | 6 | 41 |
| **Bdiv_032170** | eS10 | 0 | 134 | 15.5 | 0 | 0 |
| Bdiv_010180c | uS11 | 54 | 157 | 16.6 | 10 | 121 |
| Bdiv_040390c | uS12 | 53 | 188 | 20.8 | 3 | 65 |
| Bdiv_040260c | eS12 | 65 | 135 | 14.7 | 7 | 44 |
| Bdiv_027650 | uS13 | 34 | 154 | 17.5 | 6 | 82 |
| Bdiv_036780 | uS14 | 23 | 66 | 7.5 | 2 | 11 |
| Bdiv_001750c | uS15 | 32 | 151 | 17.4 | 6 | 41 |
| Bdiv_008760 | uS17 | 19 | 156 | 18.1 | 5 | 21 |
| **N.A.** | eS17 | 0 | 132 | 15.4 | 0 | 0 |
| Bdiv_002790 | uS19 | 44 | 149 | 17.2 | 4 | 42 |
| Bdiv_040250 | eS19 | 49 | 174 | 20.1 | 8 | 90 |
| Bdiv_033240c | eS21 | 44 | 79 | 8.7 | 4 | 29 |
| Bdiv_010070 | eS24 | 65 | 135 | 15.6 | 5 | 51 |
| Bdiv_022980c | eS25 | 22 | 104 | 11.9 | 2 | 15 |
| Bdiv_038810c | eS26 | 30 | 115 | 13.2 | 3 | 19 |
| Bdiv_036570 | eS27 | 29 | 82 | 9.1 | 2 | 26 |
| Bdiv_019090c | eS28 | 36 | 67 | 7.5 | 2 | 74 |
| Bdiv_039110c | eS30 | 72 | 61 | 6.8 | 1 | 4 |
| Bdiv_033080 | eS31 | 54 | 157 | 17.3 | 1 | 1 |
| Bdiv_010290 | RACK | 38 | 323 | 35.5 | 11 | 161 |

N.A. Ribosomal protein not annotated in *B. divergens* 1802A.

Ribosomal proteins undetected by mass spectrometry analysis are indicated in bold.

**Table S6 Ribosomal proteins identification by LC-MS/MS from the *Babesia* large subunit (related to Fig. 1)**. Sample purified from 80S ribosomes as identified by the PDB code 9YGM.

| Gene ID *B. divergens* 1802A | Unified r-prot. name | Coverage (%) | Length (amino acids) | Molecular weight (kDa) | Number of peptides | Spectral counts |
|---|---|---|---|---|---|---|
| Bdiv_038620 | uL1 | 49 | 217 | 24.4 | 8 | 138 |
| Bdiv_015470 | uL2 | 42 | 257 | 27.9 | 9 | 150 |
| Bdiv_003320c | uL3 | 43 | 395 | 44.5 | 16 | 161 |
| Bdiv_004040c | uL4 | 44 | 360 | 39.4 | 13 | 186 |
| Bdiv_027640 | uL5 | 29 | 171 | 19.7 | 5 | 58 |
| Bdiv_027090 | uL6 | 36 | 190 | 21.4 | 6 | 139 |
| Bdiv_030770c | eL6 | 43 | 212 | 23.6 | 9 | 136 |
| Bdiv_016730 | eL8 | 38 | 285 | 32.1 | 11 | 178 |
| Bdiv_028610c | uL10 | 26 | 313 | 34.0 | 9 | 97 |
| Bdiv_010230c | uL11 | 38 | 165 | 18.1 | 7 | 43 |
| Bdiv_010650 | uL13 | 33 | 202 | 23.3 | 8 | 115 |
| Bdiv_027720c | eL13 | 28 | 203 | 22.9 | 7 | 63 |
| Bdiv_035030 | uL14 | 56 | 139 | 14.8 | 7 | 79 |
| Bdiv_023300 | eL14 | 27 | 133 | 14.9 | 4 | 12 |
| Bdiv_034950 | uL15 | 20 | 147 | 16.5 | 3 | 31 |
| Bdiv_018330 | eL15 | 16 | 198 | 23.7 | 3 | 47 |
| Bdiv_016810c | uL16 | 28 | 227 | 25.7 | 6 | 37 |
| Bdiv_016720c | uL18 | 41 | 306 | 35.3 | 11 | 213 |
| Bdiv_037540 | eL18 | 18 | 247 | 27.1 | 4 | 49 |
| Bdiv_001430 | eL19 | 18 | 194 | 22.6 | 6 | 83 |
| Bdiv_037530 | eL20 | 53 | 188 | 21.6 | 10 | 126 |
| Bdiv_016780c | eL21 | 45 | 160 | 18.4 | 10 | 157 |
| Bdiv_004980c | uL22 | 18 | 194 | 22.1 | 3 | 34 |
| Bdiv_011350c | eL22 | 35 | 122 | 14.2 | 3 | 64 |
| Bdiv_039940 | uL23 | 26 | 153 | 17.3 | 4 | 33 |
| Bdiv_032900 | uL24 | 40 | 137 | 16.1 | 7 | 123 |
| Bdiv_036590c | eL24 | 19 | 199 | 22.1 | 4 | 44 |
| Bdiv_016640 | eL27 | 42 | 146 | 16.8 | 6 | 126 |
| **N.A.** | eL28 | 0 | - | - | 0 | 0 |
| Bdiv_020740c | uL29 | 37 | 123 | 14.4 | 6 | 65 |
| **Bdiv_016690** | eL29 | 0 | 59 | 6.8 | 0 | 0 |
| Bdiv_040380c | uL30 | 21 | 259 | 30.3 | 5 | 51 |
| Bdiv_034830c | eL30 | 56 | 108 | 11.9 | 4 | 43 |
| Bdiv_021180 | eL31 | 45 | 117 | 13.7 | 3 | 7 |
| Bdiv_038500 | eL32 | 24 | 132 | 15.4 | 3 | 9 |
| Bdiv_039460 | eL33 | 30 | 115 | 13.3 | 4 | 24 |
| Bdiv_019820c | eL34 | 5 | 150 | 17.0 | 1 | 2 |
| Bdiv_009850c | eL36 | 34 | 116 | 13.1 | 4 | 52 |
| Bdiv_037200 | eL37 | 7 | 98 | 10.9 | 1 | 1 |
| Bdiv_028600c | eL38 | 57 | 70 | 8.1 | 5 | 59 |
| **Bdiv_008570** | eL39 | 0 | 59 | 7.2 | 0 | 0 |
| Bdiv_024380c | eL40 | 29 | 131 | 14.9 | 3 | 12 |
| **N.A.** | eL41 | 0 | - | - | 0 | 0 |
| Bdiv_030880c | eL42 | 21 | 105 | 12.0 | 3 | 52 |
| Bdiv_029900c | eL43 | 27 | 94 | 10.6 | 2 | 71 |
| Bdiv_033480c | P1 | 45 | 117 | 12.1 | 3 | 27 |
| Bdiv_035540 | P2 | 72 | 61 | 6.5 | 2 | 22 |

N.A. Ribosomal protein not annotated in *B. divergens* 1802A.
Ribosomal proteins undetected by mass spectrometry analysis are indicated in bold.

**Table S7 Non-ribosomal protein identification by LC-MS/MS from *Babesia* (related to Fig. 1).** Sample purified from 80S ribosomes as identified by the PDB code 9YGM.

| Gene ID *B. divergens* 1802A | PiroplasmaDB protein names | Coverage (%) | Length (amino acids) | Molecular weight (kDa) | Number of Peptides | Spectral Counts |
|---|---|---|---|---|---|---|
| P02769 | serum albumin precursor [*Bos taurus*] | 31 | 607 | 69.2 | 18 | 100 |
| P13645 | Keratin, type I cytoskeletal 10 OS=*Homo sapiens* | 28 | 593 | 59.5 | 12 | 29 |
| P35908 | Keratin, type II cytoskeletal 2 epidermal OS=Homo sapiens | 27 | 639 | 65.4 | 12 | 21 |
| Bdiv_010550 | splicing factor, arginine/serine-rich 3 | 24 | 235 | 27.5 | 5 | 15 |
| Bdiv_027340 | ATP synthase F1 beta chain | 23 | 515 | 55.4 | 7 | 12 |
| Bdiv_031040 | vesicle-associated membrane protein | 22 | 225 | 25.2 | 3 | 6 |
| P08515 | Glutathione S-transferase class-mu 26 kDa isozyme OS=*Schistosoma japonicum* | 22 | 218 | 25.5 | 6 | 19 |
| Bdiv_031930 | NAC domain containing protein | 21 | 208 | 22.3 | 3 | 5 |
| P00761 | Trypsin precursor - sus scrofa | 20 | 741 | 79.4 | 5 | 15 |
| Bdiv_016790c | NAC domain containing protein | 19 | 156 | 16.7 | 2 | 6 |
| Bdiv_037120c | hsp90 protein | 15 | 708 | 81.6 | 7 | 19 |
| P04264 | Keratin, type II cytoskeletal 1 OS=*Homo sapiens* | 15 | 644 | 66.0 | 8 | 19 |
| P35527 | Keratin, type I cytoskeletal 9 OS=*Homo sapiens* | 15 | 623 | 62.0 | 6 | 10 |
| Bdiv_001940 | aspartate transcarbamoylase | 14 | 382 | 42.9 | 4 | 10 |
| Bdiv_007280* | RNA chaperone ProQ; FinO, | 14 | 115 | 12.1 | 1 | 1 |

| | | | | | | |
|---|---|---|---|---|---|---|
| | [*Escherichia coli* (strain K12)] | | | | | |
| Bdiv_037240c | nucleolar GTP-binding protein 1 | 13 | 590 | 67.5 | 6 | 12 |
| Bdiv_015690c | 50 kDa surface antigen | 13 | 445 | 47.9 | 5 | 9 |
| Bdiv_033760 | Apicomplexa-specific secreted signal peptide [*Babesia ovis*] | 13 | 250 | 27.9 | 2 | 2 |
| P81605 | Dermcidin precursor - *Homo sapiens* | 13 | 110 | 11.3 | 2 | 2 |
| Bdiv_039420 | BOP1NT (NUC169) domain containing protein | 12 | 709 | 81.3 | 7 | 43 |
| Bdiv_014120c | mRNA turnover protein 4 mrto4 like protein [*Babesia gibsoni*] | 12 | 225 | 25.2 | 2 | 8 |
| Bdiv_007890 | actin | 12 | 376 | 41.9 | 3 | 9 |
| Bdiv_004780 | transmembrane protein, putative [*Babesia ovis*] | 11 | 234 | 27.6 | 2 | 2 |
| Bdiv_031450 | glycosylphosphatidylinositol-anchored merozoite surface protein | 11 | 324 | 35.0 | 2 | 6 |
| Bdiv_011020 | gar1 protein RNA binding region containing protein | 10 | 191 | 20.0 | 1 | 1 |
| Bdiv_009840c* | pre-60S ribosome [*Saccharomyces cerevisiae* S288C] | 9 | 161 | 18.4 | 1 | 1 |
| Bdiv_009750 | pescadillo N-terminus family protein | 8 | 468 | 53.2 | 3 | 14 |
| Bdiv_027520c | proliferation-associated protein 2g4 | 8 | 387 | 42.0 | 3 | 10 |
| Bdiv_015840c | apolipoprotein N-acyltransferase [*Babesia caballi*] | 8 | 144 | 15.3 | 1 | 1 |
| Bdiv_009230c | p29 protein | 8 | 201 | 23.5 | 1 | 1 |

| | | | | | | |
|---|---|---|---|---|---|---|
| Bdiv_017280c | adenine nucleotide translocase | 7 | 299 | 33.4 | 2 | 6 |
| Bdiv_002420 | gliding-associated protein 45 | 7 | 184 | 21.4 | 1 | 2 |
| Bdiv_032380 | WD domain, G-beta repeat containing protein | 7 | 439 | 48.4 | 3 | 6 |
| Bdiv_025320c | transporter | 7 | 467 | 50.7 | 3 | 6 |
| Bdiv_035860 | Translocation protein [Babesia sp. Xinjiang] | 7 | 404 | 46.6 | 2 | 2 |
| Bdiv_032250 | proteasome A-type and B-type family protein | 7 | 257 | 28.3 | 1 | 1 |
| Bdiv_026360c | hypothetical protein, conserved | 7 | 172 | 18.1 | 1 | 1 |
| Bdiv_023990c | apical membrane antigen 1 | 6 | 605 | 67.5 | 3 | 8 |
| Bdiv_002930c | 60S ribosomal protein L7-like 1 | 6 | 234 | 26.5 | 1 | 2 |
| Bdiv_018040c | proteasome A-type and B-type family protein | 6 | 252 | 27.5 | 1 | 1 |
| Bdiv_038840c | secreted antigen 1 | 6 | 533 | 57.7 | 2 | 2 |
| Bdiv_032480 | vacuolar protein sorting 29 | 6 | 246 | 27.0 | 1 | 1 |
| Bdiv_021150c | Ser/Arg-rich splicing factor | 6 | 199 | 24.0 | 1 | 3 |
| Bdiv_005420 | heat shock protein 60 | 6 | 556 | 59.7 | 2 | 6 |
| Bdiv_011500 | cytochrome c oxidase subunit ApiCOX25, putative [*Plasmodium yoelii*] | 6 | 254 | 29.3 | 1 | 1 |
| P00766 | Chymotrypsinogen A - *Bos taurus* | 6 | 245 | 25.7 | 1 | 1 |
| Bdiv_017140 | COPI associated protein, putative [*Babesia caballi*] | 6 | 276 | 31.2 | 1 | 1 |
| Bdiv_014380 | secreted antigen 1 | 6 | 313 | 34.8 | 1 | 1 |
| Bdiv_020050c | major facilitator superfamily MFS-1 [*Babesia ovata*] | 5 | 486 | 53.3 | 2 | 8 |

| | | | | | | |
|---|---|---|---|---|---|---|
| Bdiv_018730 | RNA processing factor 1 | 5 | 301 | 35.2 | 1 | 7 |
| Bdiv_007850 | secreted antigen 1 | 5 | 497 | 54.5 | 2 | 2 |
| Bdiv_036170c | fatty acyl-CoA synthetase family protein | 5 | 673 | 74.7 | 2 | 2 |
| Bdiv_011410c | cytochrome oxidase subunit I, putative [*Babesia ovata*] | 5 | 311 | 34.2 | 1 | 2 |
| Bdiv_040150 | Mak16 family protein | 5 | 222 | 26.2 | 1 | 2 |
| Bdiv_019270 | proliferating cell nuclear antigen 1 | 5 | 277 | 30.8 | 1 | 1 |
| Bdiv_031730c | membrane protein | 5 | 367 | 40.5 | 1 | 1 |
| Bdiv_025500c | erythrocyte binding protein 37.2 precursor | 5 | 316 | 34.7 | 1 | 1 |
| Bdiv_003280 | Brix domain containing protein | 4 | 323 | 36.5 | 1 | 1 |
| Bdiv_009700 | Ribosome biogenesis protein nsa1; 60S [*Schizosaccharomyces*] | 4 | 590 | 66.1 | 2 | 3 |
| Bdiv_036030 | Pre mRNA splicing factor CWF19, cell cycle control protein, putative [*Babesia bigemina*] | 4 | 310 | 35.6 | 1 | 1 |
| Bdiv_006980c | hypothetical protein, conserved | 4 | 241 | 25.8 | 1 | 1 |
| Bdiv_033560c | enolase (2-phosphoglycerate dehydratase) | 4 | 442 | 48.0 | 1 | 2 |
| Bdiv_013400c | secreted antigen 1 | 4 | 567 | 62.3 | 1 | 2 |
| Bdiv_025200c | aspartyl aminopeptidase | 4 | 447 | 49.3 | 1 | 2 |
| 2497269 | Keratin, type i cytoskeletal 12 [*Homo sapiens*] | 4 | 494 | 53.5 | 1 | 1 |
| Bdiv_036250c | 2-methylcitrate synthase [*Babesia caballi*] | 4 | 170 | 19.0 | 1 | 1 |

| | | | | | | |
|---|---|---|---|---|---|---|
| Bdiv_039450c | eukaryotic porin protein [*Babesia caballi*] | 4 | 298 | 32.8 | 1 | 1 |
| Bdiv_023170 | glutamate dehydrogenase | 3 | 455 | 50.6 | 1 | 1 |
| Bdiv_003860c | rhomboid 4 | 3 | 632 | 69.8 | 1 | 1 |
| Bdiv_003340 | Clathrin interactor EPSIN 1 [*Babesia sp. Xinjiang*] | 3 | 462 | 51.9 | 1 | 2 |
| Bdiv_011370c | PCI domain containing protein | 3 | 431 | 48.9 | 1 | 2 |
| Bdiv_012730c | CAAX metallo endopeptidase | 3 | 448 | 52.1 | 1 | 2 |
| Bdiv_036160c | secreted antigen 1 | 3 | 566 | 62.1 | 1 | 2 |
| Bdiv_022170c | nonsense-mediated mRNA decay protein | 3 | 493 | 56.5 | 1 | 1 |
| Bdiv_005070 | nucleolar GTP-binding protein 2 | 3 | 636 | 71.9 | 1 | 1 |
| Bdiv_012850 | putative protein phosphatase 2C 8 [*Babesia sp. Xinjiang*] | 3 | 393 | 45.2 | 1 | 1 |
| Bdiv_025360c | transcription or splicing factor-like protein | 3 | 483 | 54.1 | 1 | 1 |
| P08659 | Luciferin 4-monooxygenase Firefly luciferase OS=*Photinus pyralis* | 3 | 550 | 60.7 | 1 | 1 |
| Bdiv_019870 | GTP binding protein | 3 | 423 | 46.8 | 1 | 1 |
| Bdiv_003940 | sulfate transporter | 3 | 723 | 80.1 | 1 | 1 |
| Bdiv_003010c | secreted antigen 1 | 3 | 298 | 33.4 | 1 | 1 |
| Bdiv_011800 | membrane protein | 2 | 716 | 80.4 | 1 | 2 |
| Bdiv_002620 | myb-like DNA-binding domain containing protein | 2 | 537 | 61.5 | 1 | 1 |
| Bdiv_036530 | ATP synthase [*Babesia duncani*] | 2 | 544 | 62.7 | 1 | 2 |
| Bdiv_018060* | Mitochondrial RNA binding complex 1 | 2 | 643 | 71.3 | 1 | 2 |

| | | | | | | |
|---|---|---|---|---|---|---|
| | subunit [*Trypanosoma brucei*] | | | | | |
| Bdiv_036610c | carbamoyl phosphate synthetase | 2 | 1630 | 179.8 | 2 | 2 |
| Q7Z794 | Keratin, type II cytoskeletal 1b OS=*Homo sapiens* | 2 | 576 | 61.7 | 1 | 2 |
| Bdiv_011530 | RIBONUCLEASE Z domain containing protein, putative | 2 | 661 | 74.1 | 1 | 1 |
| Bdiv_014590c | Phosphatidate cytidylyltransferase domain-containing protein | 2 | 543 | 59.9 | 1 | 2 |
| P13647 | Keratin, type II cytoskeletal 5 OS=*Homo sapiens* | 2 | 590 | 62.3 | 1 | 1 |
| Bdiv_024840 | WD repeat domain containing protein | 2 | 848 | 94.6 | 1 | 1 |
| Bdiv_034280 | DNA repair protein (mre11) family protein | 2 | 967 | 108.0 | 1 | 1 |
| Bdiv_034410c* | mL113 Mitochondria [*Chlamydomonas reinhardtii*] | 2 | 597 | 67.4 | 1 | 1 |
| 547751 | Keratin, type i cytoskeletal 17 (cytokeratin 17) [*Homo sapiens*] | 2 | 432 | 48.1 | 1 | 1 |
| Bdiv_010860 | micro-fibrillar-associated protein 1 C-terminus containing protein | 2 | 450 | 51.9 | 1 | 1 |
| Bdiv_010090 | eukaryotic translation initiation factor 3 subunit | 2 | 732 | 85.2 | 1 | 1 |
| Bdiv_028060c | translation initiation factor 3 subunit 10 | 2 | 990 | 117.0 | 1 | 1 |
| Bdiv_031890 | kynurenine 3-monooxygenase and-related flavo monooxygenas | 1 | 1163 | 129.6 | 1 | 10 |

| | | | | | | |
|---|---|---|---|---|---|---|
| | e family protein, putative [*Babesia ovata*] | | | | | |
| Bdiv_003100 | helicase | 1 | 1920 | 217.0 | 1 | 1 |
| Bdiv_012610c | ubiquitin thioesterase [*Babesia gibsoni*] | 1 | 4400 | 497.2 | 3 | 3 |
| Bdiv_012920 | AP2 domain transcription factor AP2X-11 [Babesia caballi] | 1 | 629 | 68.4 | 1 | 3 |
| Bdiv_030760 | ATP-dependent metalloprotease FtsH family protein | 1 | 705 | 78.4 | 1 | 1 |
| Bdiv_010010c | RNA polymerase I specific transcription initiation factor | 1 | 1030 | 117.3 | 1 | 1 |
| Bdiv_020780c | vacuolar sorting-associated protein 36 [*Babesia ovis*] | 1 | 468 | 53.3 | 1 | 1 |
| Bdiv_014910 | exportin 1 | 1 | 1189 | 138.1 | 1 | 2 |
| Bdiv_014470c | ubiquitin-activating enzyme E1 | 1 | 1010 | 113.7 | 1 | 1 |

* Protein name/identification resulting from Blast using HHpred interactive server for protein homology detection and structure prediction.

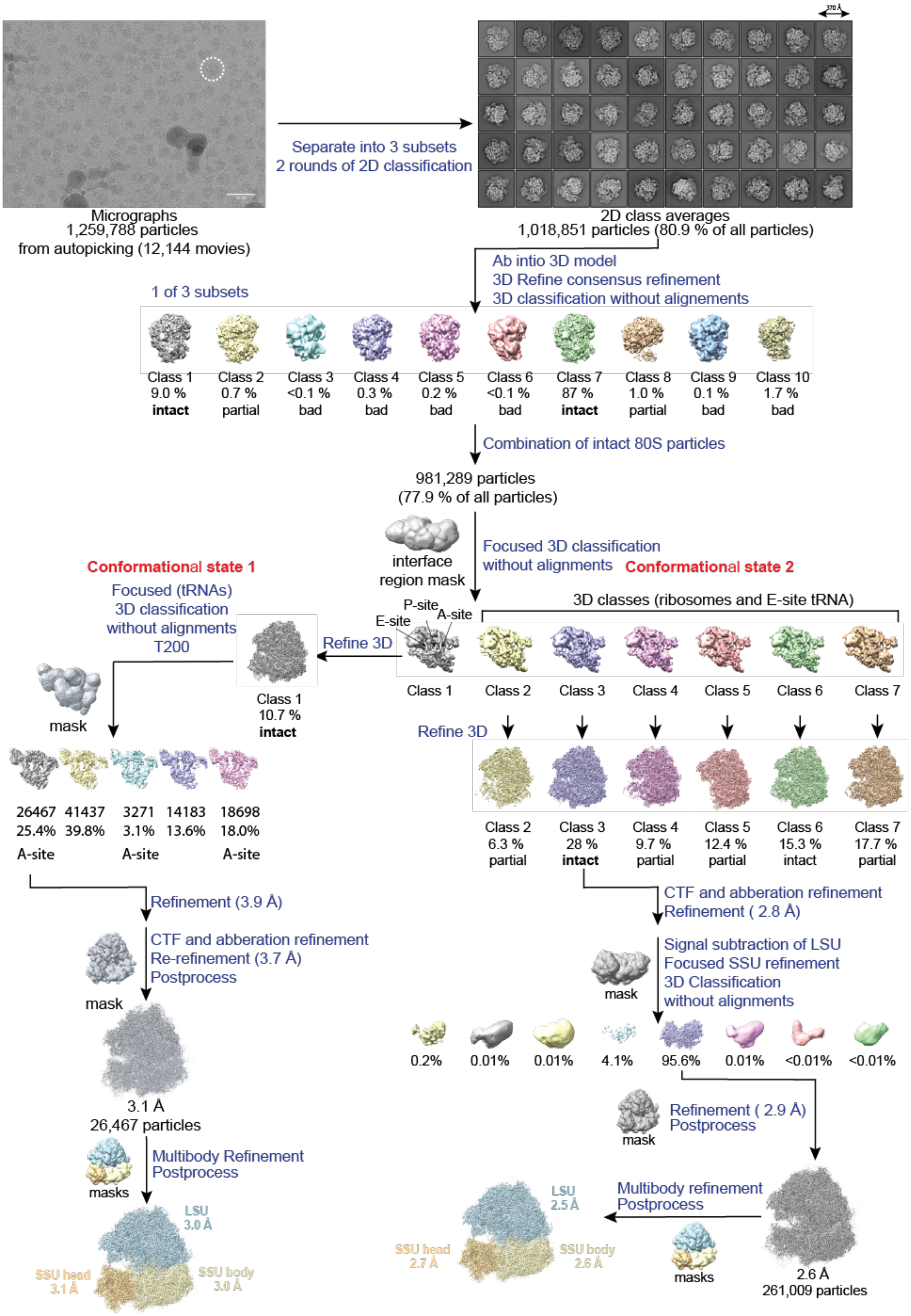


**Fig. S1 Data collection and data processing scheme for the structure of *Babesia* ribosomes by single-particle cryo-EM (related to Fig. 1).** A typical drift-corrected cryo-EM micrograph of purified ribosomes is shown (a white dotted circle shows a representative particle). 2D class averages show densities corresponding to the large (LSU) and the small subunits (SSU). The flowchart of the 3D classification illustrates the processing scheme used. The data was split into three subsets before 2D classification, where one of the three subsets is shown. We combined intact particles for further processing (see *Methods*).

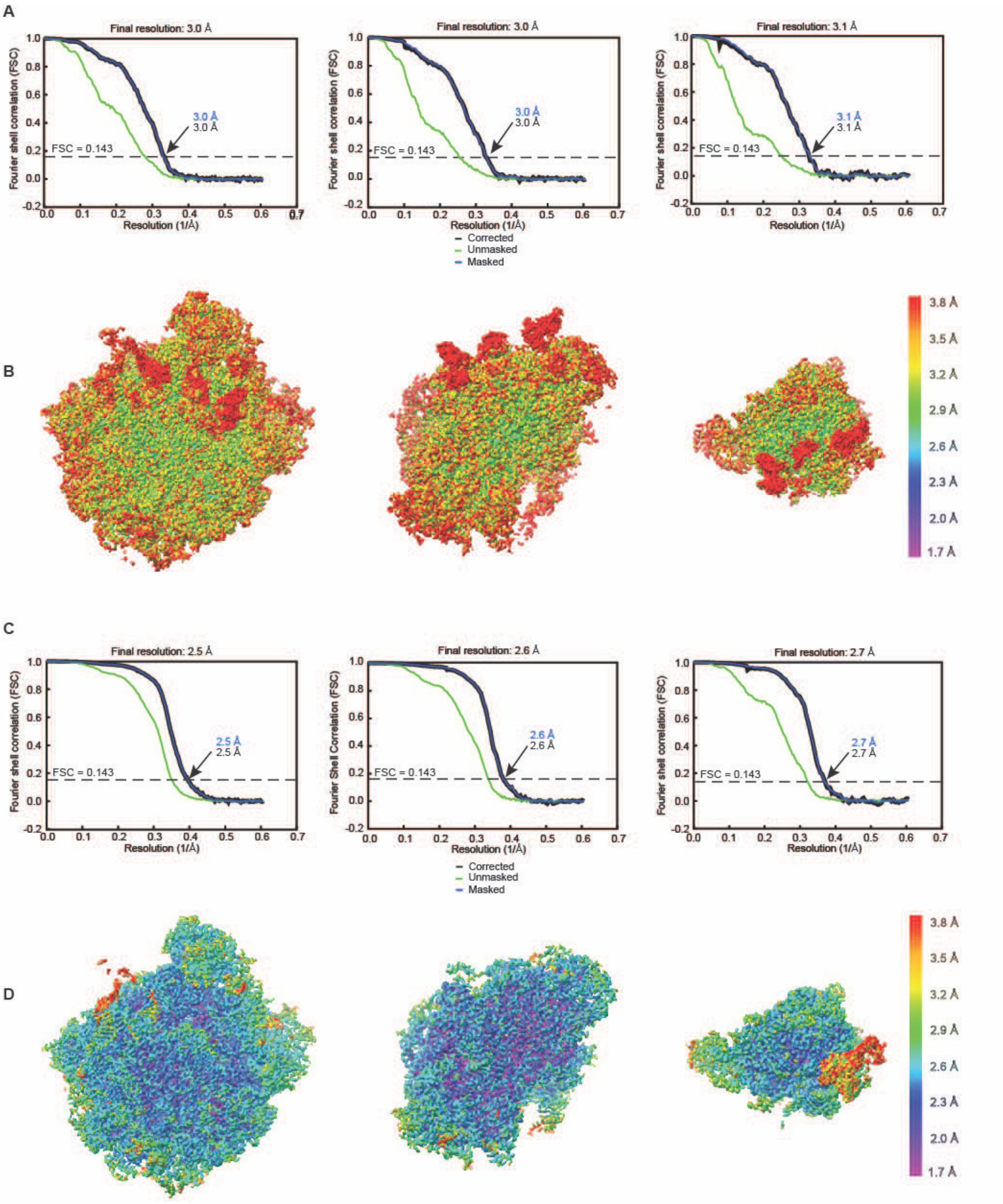


**Fig. S2 Resolution estimation of ribosome subdomains after multi-body refinement (related to Fig. 1). A** 3DC1 fourier shell correlation (FSC) curves as a function of spatial frequency showing a nominal resolution relative to the gold-standard FSC = 0.143 criterion of 3.0, 3.0, and 3.1 Å for the LSU, the body of the SSU, and the head of the SSU, respectively. Density is visible for tRNAs occupying the E, P, and A sites. **B** Estimated local resolution map of the interface view of the LSU, SSU body, and of the SSU head, respectively. **C** 3DC3 fourier shell correlation (FSC) curves as a function of spatial frequency showing a nominal resolution relative to the gold-standard FSC = 0.143 criterion of 2.5, 2.6, and 2.7 Å for the LSU with an E-site tRNA density, the body of the SSU, and the head of the SSU with an E-site density, respectively. **D** Estimated local resolution map of the interface view of the LSU and an E-site tRNA, SSU body, and of the SSU head and E-site tRNA, respectively.

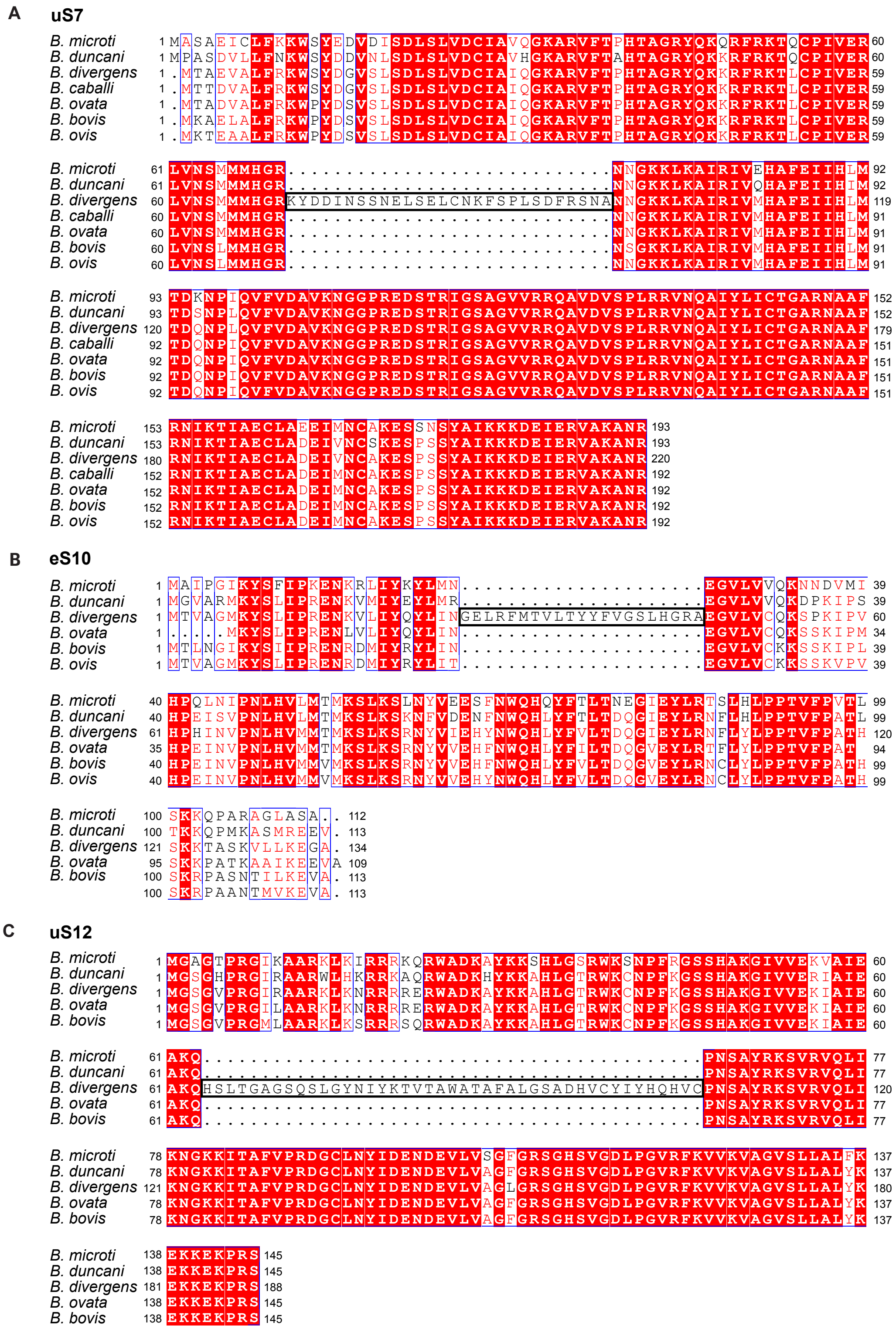


**Fig. S3 Sequence alignment of ribosomal proteins from the small subunit between *Babesia* species based on genome annotation from PiroplasmaDB (related to Fig. 1).** Sequence alignment was generated using Clustal Omega and is displayed using ESPript3. White letters on a red background correspond to identical residues, while red letters on a white background indicate conservation. A black box indicates a portion of a sequence that was misannotated in *B. divergens* 1802A strain, as it overlaps with an intron in *B. divergens* Rouen strain, consistent with the other *Babesia* strains displayed in the alignment. The numbering is based on the *B. microti* sequence. **A** uS7 sequence alignment. PiroplasmaDB gene IDs: *B. microti*: BmR1_04g08092, *B. duncani*: BdWA1_002425, *B. divergens* (1802A): Bdiv_017820c, *B. caballi*: BcabD6B2_13830, *B. ovata*: BOVATA_042040, *B. bovis*: BBOV_I000230, and *B. ovis*:BaOVIS_000820. **B** eS10 sequence alignment. *B. microti*: BmR1_04g06575, *B. duncani*: BdWA1_001572, *B. divergens* (1802A): Bdiv_032170, *B. ovata*: BOVATA_000860, *B. bovis*: BBOV_III009270, and *B. ovis*: BaOVIS_009200. **C** uS12 sequence alignment. *B. microti*: BMR1_02g01585, *B. duncani*: BdWA1_002523, *B. divergens* (1802A): Bdiv_040390c, *B. ovata*: BOVATA_045610, and *B. bovis*: BBOV_III000970.

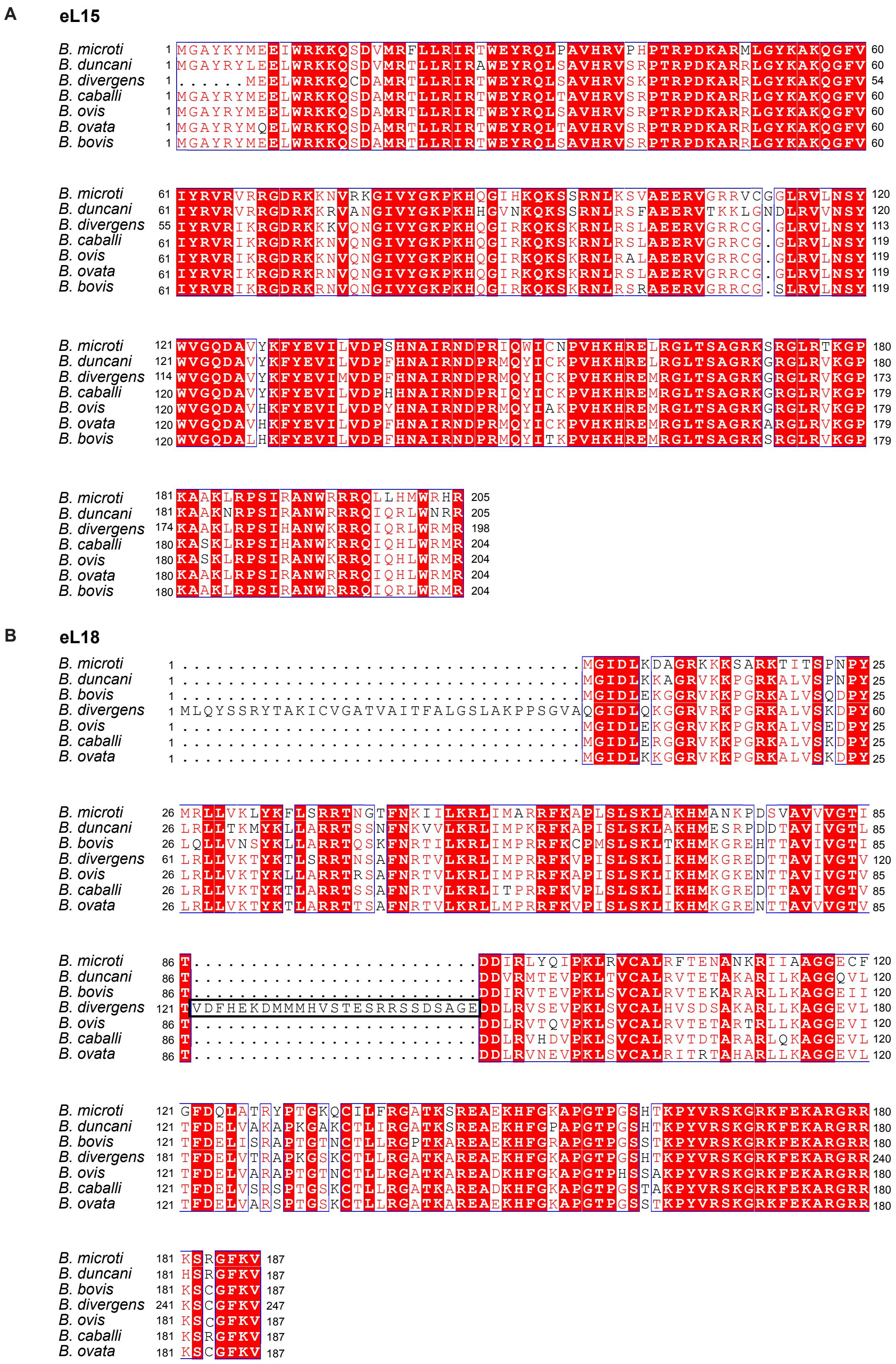
A
eL15
B. microti
B. duncani
B. divergens
B. caballi
B. ovis
B. ovata
B. bovis
B
eL18

**C** **eL24**

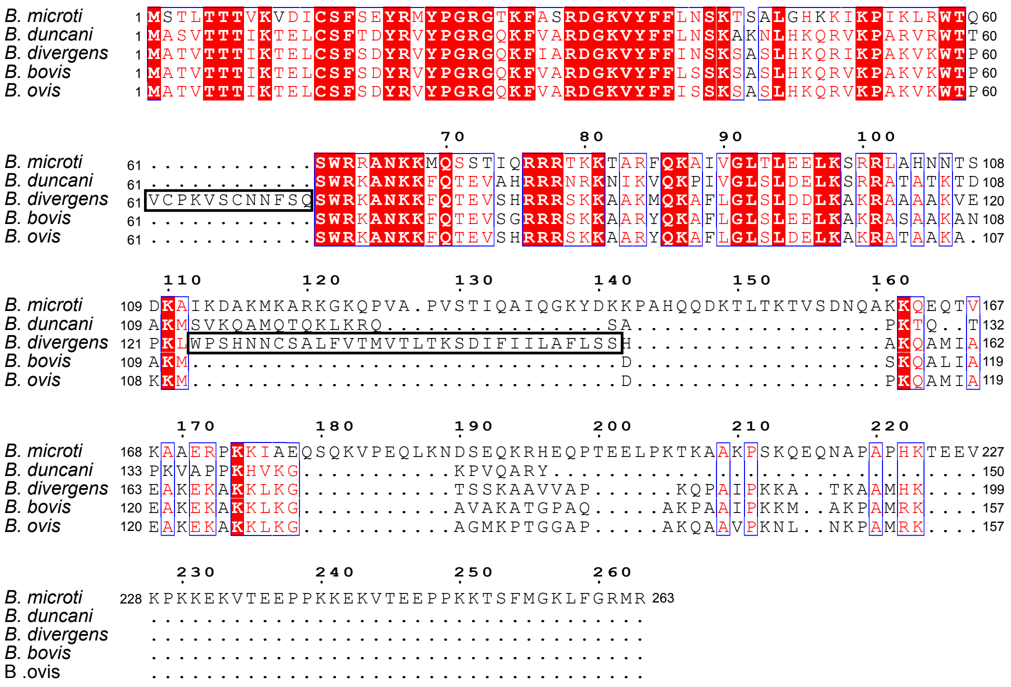


**Fig. S4. Sequence alignment of ribosomal proteins from the large subunit between *Babesia* species based on genome annotation from PiroplasmaDB (related to Fig. 1).** Sequence alignment was generated and displayed as described in **Fig. S3**. **A** eL15 sequence alignment. Based on cryo-EM density analysis,the eL15 sequence was misannotated and contains the N-terminal sequence (MGAYRY) found in all other *Babesia* species included in the sequence alignment. PiroplasmaDB gene IDs: *B. microti*: BmR1_04g09685, *B. duncani*: BdWA1_002482, *B. divergens* (1802A): Bdiv_018330, *B. caballi*: BcabD6B2_14580, *B. ovis*: BaOVIS_001320, *B. ovata*: BOVATA_042510, and *B. bovis*: BBOV_III000550. **B** eL18 sequence alignment. PiroplasmaDB gene IDs: *B. microti*: BmR1_04g04880, *B. duncani*: BdWA1_000922, *B. divergens* (1802A): Bdiv_037540, *B. caballi*: BcabD6B2_03030, *B. ovata*: BOVATA_006680, *B. bovis*: BBOV_III004640, and *B. ovis*:BaOVIS_004890. **C** eL24 sequence alignment. *B. microti*: BMR1_01G02266, *B. duncani*: BdWA1_002225, *B. divergens* (1802A): Bdiv_036590c, *B. bovis*: BBOV_III003570, and *B. ovis*: BaOVIS_004000.

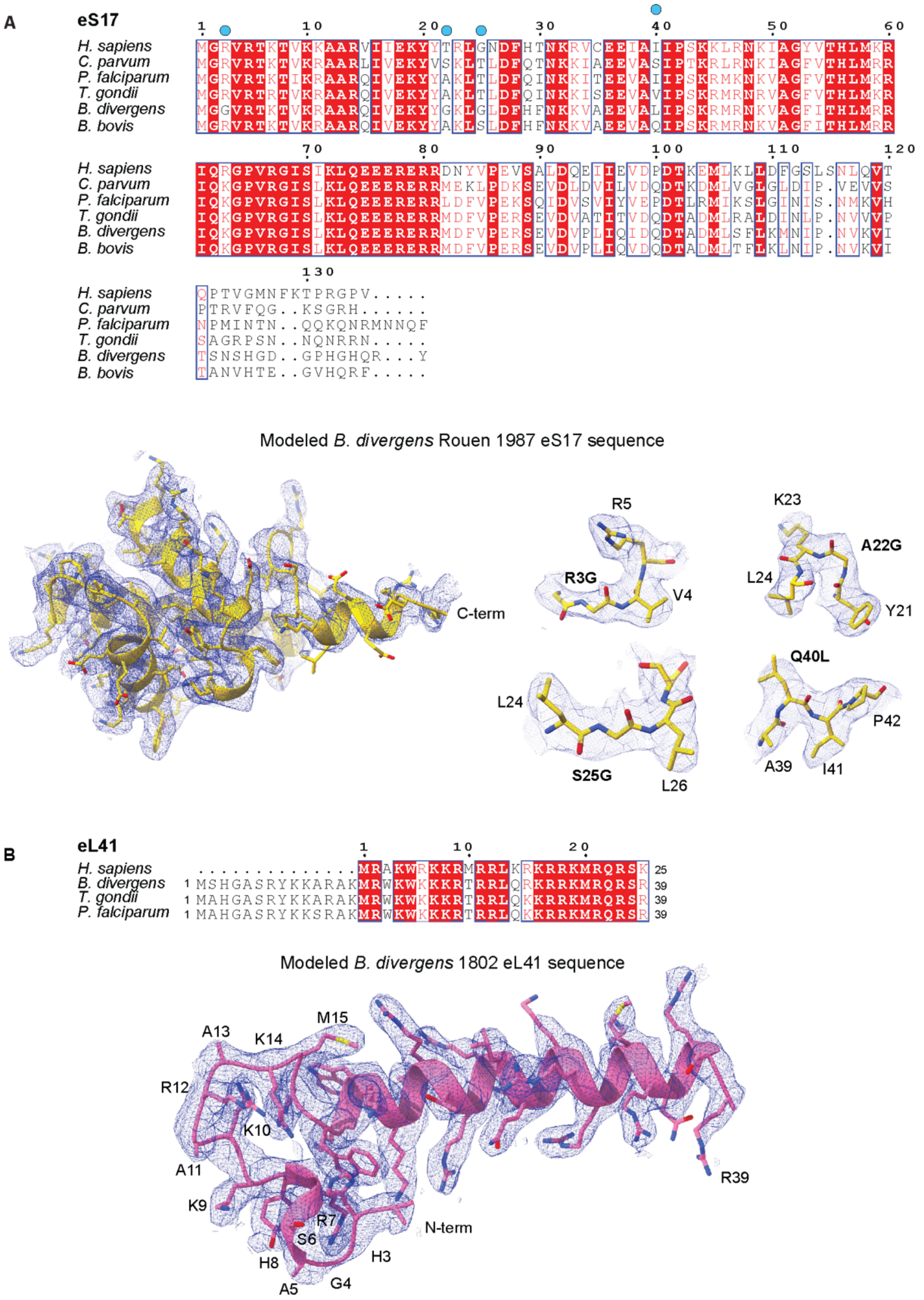


**Fig. S5 Model fitting of eS17 and eL41 into the *Babesia* ribosome density (related to Figs. 1 and 3).** The sequence alignment was generated and displayed as described in **Fig. S3**. The numbering is based on the human sequence. **A** The top panel shows the eS17 sequence alignment. Cyan circles indicate residue diffrences between *B. divergens* and *B. bovis*. *H. sapiens*: P08708, *C. parvum*: cgd5_3720, *P. falciparum*: PF3D7_1242700, *T. gondii*: TGME49_20784, *B. divergens* (Rouen 1987): CCSG02000001.1:1890202..1890772 (supercontig), *B. bovis*: BBOV_III001250. The bottom panel shows the atomic model of eS17. eS17 *B. bovis* sequence (BBOV_III001250) used for initial model building (Alphafold2) and further mutated over four residues (R3G, A22G, S25G, Q40L) to match the corresponding *B. divergens* protein sequence identified from the Rouen 1987 strain. The left panel displays the complete eS17 model (threshold level of 5.0), while the right panel highlights the mutated residues from *B. bovis* to *B. divergens* (threshold level of 6.0). **B** The top panel shows the eL41 sequence alignment. eL41 gene IDs. *H. sapiens* : P62945c, *B. divergens* (1802): JAHBMH010000034: 30674..30988 supercontig, *T. gondii*: TGME49_213580, and *P. falciparum*: PF3D7_1144300. The bottom panel shows the atomic model of eL41 in the density map (threshold 5.0), including the N-terminal extension (residues 2–14) found in apicomplexan parasites.

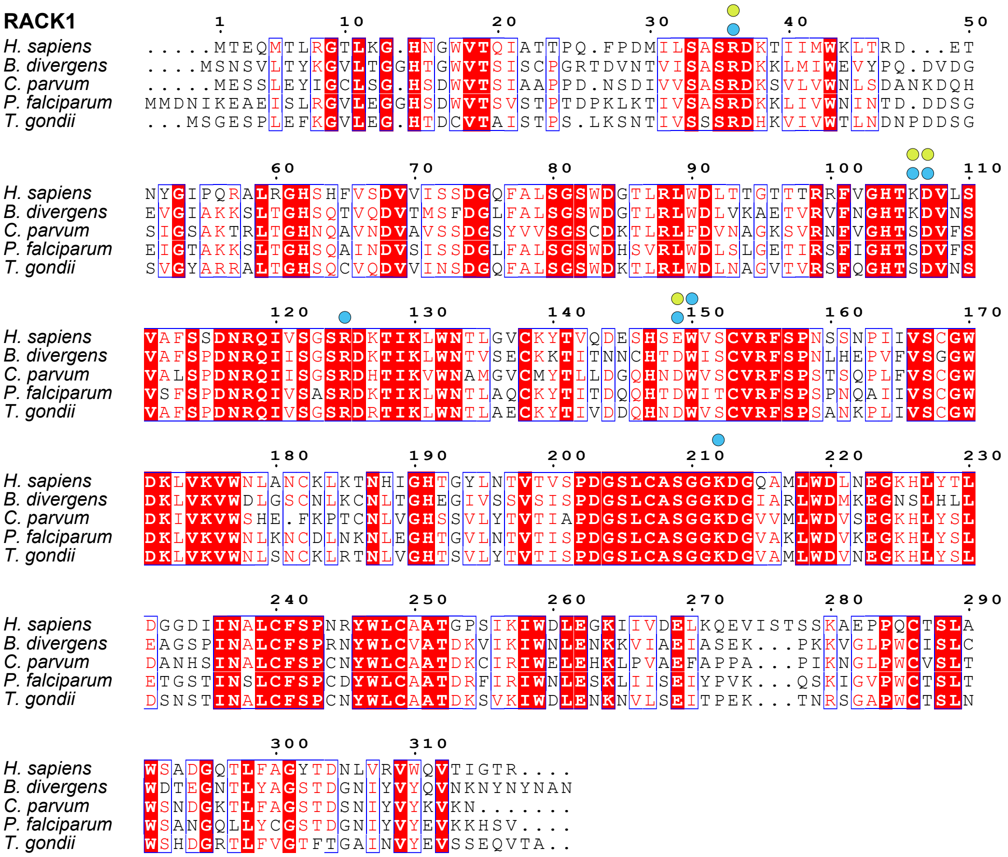

**Fig. S6 Sequence alignment of RACK1 (related to Figs. 1 and 3).** The sequence alignment was generated and displayed as described in **Fig. S3**. The numbering is based on the human sequence. Cyan and green circles indicate RACK1 residues interacting with eS17 through hydrogen bonds or non-covalent interactions in *B. divergens* and humans, respectively. VEuPathDB gene IDs: *H. sapiens*: P63244, *B. divergens*: Bdiv_010290, *C. parvum*: cgd2_1870, *P. falciparum*: PF3D7_0826700, and *T. gondii*: TGME49_216880.

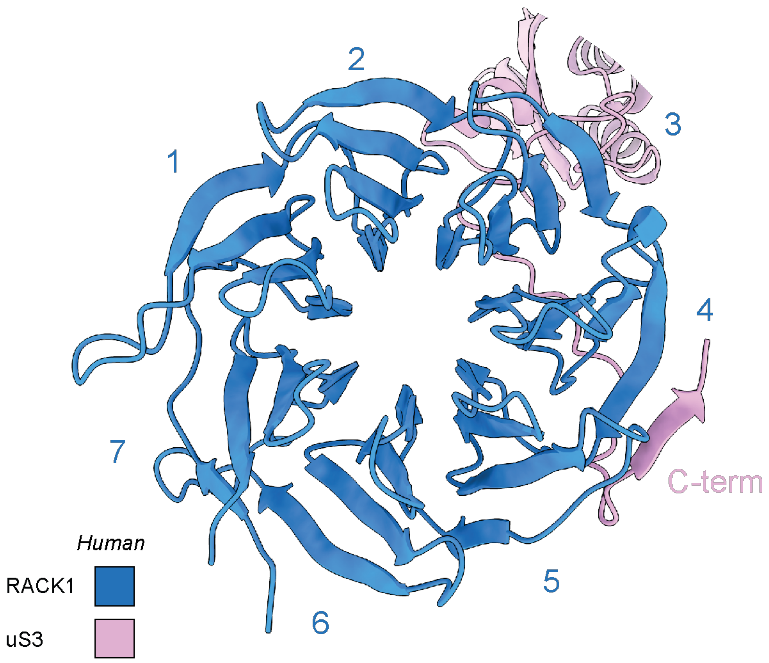


**Fig. S7 Overview of the localization of the human C-terminal tail of uS3 relative to human RACK1 within the head of the SSU (PDB 8QOI, related to Fig. 3)**. View from the top of RACK1, showing each propeller blade (1-6, blue). The C-terminal (C-term) tail of uS3 interacts with RACK1 (pink) between the propeller blades 4 and 5.
